# A reparative neutrophil subpopulation promotes spinal cord regeneration in zebrafish by controlling macrophage inflammation via Il-4

**DOI:** 10.64898/2026.02.25.707887

**Authors:** Xiaobo Tian (田晓波), Alberto Docampo-Seara, Kim Heilemann, Friederike Kessel, Daniela Zöller, Anja Bretschneider, Thomas Becker, Catherina G. Becker

## Abstract

In mammals, a dysregulated immune response is detrimental to spinal cord repair. In zebrafish, which are capable of spinal cord regeneration, the immune response promotes regeneration. Neutrophils are the first immune cells to arrive at a spinal cord injury site, but their role in successful regeneration is not fully understood. Here we show that ablating neutrophils, including a subpopulation that expresses the cytokine *il4*, increases expression of *il1b* (coding for Il-1β) in macrophages/microglia and impairs anatomical and functional recovery after a spinal cord injury in larval zebrafish. Regeneration is fully rescued by over-expression of *il4* alone or experimentally reducing Il-1β levels. Disruption of *il4* mimics the detrimental effect of neutrophil ablation for axonal regeneration and is also rescued by reducing Il-1β levels. Hence, after spinal cord injury, a pro-regenerative neutrophil subpopulation promotes spinal cord regeneration in larval zebrafish by controlling expression of *il1b* in macrophages/microglia. For this neutrophil action, *il4* expression is necessary and sufficient.

**HIGHLIGHTS:**

- Neutrophil ablation impairs spinal cord repair in zebrafish
- The neutrophil response can be replaced by reducing Il-1β levels
- A pro-regenerative subpopulation of neutrophils expresses *il4*
- *il4* overexpression fully rescues effects of neutrophil ablation

## INTRODUCTION

Spinal cord injury leads to damage of axonal tracts that is not repaired in most mammals, including humans, often leading to permanent disability (Courtine & Sofroniew, 2019; Fehlings *et al*, 2017). In contrast, larval and adult zebrafish regrow axons and recover locomotion after complete spinal cord transection (Becker & Becker, 2014; Tendolkar & Mokalled, 2025; Tsata & Wehner, 2021). The immune reaction after injury in mammals is protracted, causing an inflammatory environment that is not conducive to regeneration (Greenhalgh *et al*, 2020; Rodgers *et al*, 2022; Stewart *et al*, 2025). Neutrophils and macrophages/microglia are the major cell types of the innate immune response that persist at the injury site in mammals (Beck *et al*, 2010).

In both, regenerating and non-regenerating systems, neutrophils are the first immune cells that arrive at a spinal cord injury site (Becker & Becker, 2022; Galli *et al*, 2011). Neutrophils negatively influence the balance of pro- and anti-inflammatory cytokines expressed by macrophages and are generally found to be detrimental for repair (Diao *et al*, 2025; Dolma & Kumar, 2021; Kitade *et al*, 2023; Kumar *et al*, 2018; Saiwai *et al*, 2010). However, a sub-population of neutrophils has recently been discovered that promotes regeneration of the optic nerve in mice by secreting several growth factors (Sas *et al*, 2020). Understanding neutrophil phenotypes is crucial also for therapeutic approaches. For example, transplantation of neural stem cells to repair the spinal cord is aided by removal of neutrophils (Nguyen *et al*, 2017). Influencing the neutrophil phenotype towards a more pro-regenerative one or preserving pro-regenerative neutrophils could aid transplantation experiments.

In the highly accessible larval zebrafish model of spinal cord injury (Alper & Dorsky, 2022), neutrophils also exhibit a strong accumulation in a spinal cord injury site within minutes after spinal cord injury (de Sena-Tomas *et al*, 2024; John *et al*, 2025; Tsarouchas *et al*, 2018). Preventing neutrophils from reaching the injury site or accelerating neutrophil resolution improves aspects of regeneration, suggesting negative roles for neutrophils also in regenerating zebrafish (de Sena-Tomas *et al*., 2024). However, neutrophils may favourably influence the mechanical properties of the lesion tissue (John *et al*., 2025). In an experimental situation in which macrophages are missing, the increased number of remaining neutrophils contribute to an increased expression of the pro-inflammatory cytokine *il1b* (Tsarouchas *et al*., 2018). High levels of *il1b* expression inhibit spinal cord regeneration in larval zebrafish (Oprişoreanu *et al*, 2023). Removing the neutrophils partially rescues reduced axon regrowth in the absence of macrophages, indicating that macrophages can suppress an anti-regenerative phenotype in neutrophils (Tsarouchas *et al*., 2018). However, whether subpopulations of neutrophils exist that may be pro- or anti-regenerative and how these may signal to other cells in the injury site is not known.

Here we characterize a substantial sub-population of neutrophils that expresses the cytokine *il4* and supports regeneration. Ablating neutrophils impairs regeneration in larval zebrafish and leads to upregulation of *il1b* in macrophages. Reducing elevated levels of Il-1β or over-expression of *il4* fully rescues regeneration after pan-neutrophil ablation. Thus, we identify a pro-regenerative sub-population of neutrophils that promotes regeneration by reducing *il1b* expression in macrophages/microglia via Il-4.

## RESULTS

### Selective ablation of neutrophils impairs spinal cord regeneration

Neutrophils arrive in the injury site within minutes, leading to an early peak of neutrophil invasion of the injury site at 4 hours post-lesion (hpl) (Tsarouchas *et al*., 2018). To establish a role of neutrophils in spinal cord regeneration in zebrafish, we decided to selectively remove these cells prior to injury and thus reduce neutrophil invasion of the injury site. To that aim, we used transgenic fish in which a fusion protein of the bacterial enzyme nitroreductase (NTR) and the fluorescent reporter mCherry was expressed under the control of the regulatory sequences of the neutrophil-specific gene *mpx* (*mpx:NTR-mCherry*; see Material and Methods). Animals were pre-incubated with the prodrug Metronidazole (MTZ), which is turned into a cytotoxic compound in cells expressing NTR, 8 hours prior to the time point of spinal cord injury at 3 days post-fertilization (dpf). To monitor the effectiveness of this treatment, we determined cell death in the caudal hematopoietic tissue (CHT), which is ventral to the prospective injury site and the major place of origin for immune cells that invade the lesion site (Tsarouchas *et al*., 2018). We found co-labelling of mCherry and the cell death indicator acridine orange in the CHT in 89.7% of all mCherry-expressing cells in unlesioned animals at 3 dpf, indicating that these were dead cells (Suppl. Fig. 1A-C). Double-labelling of macrophages with an anti-Mfap4 antibody indicated that the number of macrophages in the CHT was not changed by neutrophil ablation and that some macrophages had engulfed mCherry-positive profiles, providing further evidence for successful ablation (Suppl. Fig. 1D-F). Consequently, at 4 hpl, *mpx:NTR-mCherry*-positive neutrophils accumulated in the lesion site in control animals, but in MTZ-treated animals this was not the case. The sparse mCherry signal in the lesion site was associated only with macrophages (Suppl. Fig. 1G-I). We confirmed successful reduction of neutrophil numbers in the lesion using an antibody to Mpx (Suppl. Fig. 1J, K).

For a time-course analysis, we crossed *mpx:NTR-mCherry* into a second neutrophil reporter line (*mpx:gfp)* to avoid confounding debris. In control animals that had not been incubated with MTZ, we observed a sharp peak in neutrophil abundance in the injury site at 4 hours in accordance with previous observations (Tsarouchas *et al*., 2018). In contrast, in experimental animals, this peak was almost completely abolished. Low numbers of neutrophils were present in experimental animals for up to 24 hpl, at which time there was no difference to the low numbers of neutrophils in control animals anymore (Fig. 1A-C). Hence, we were able to abolish the peak of neutrophil invasion.

**Figure 1.**
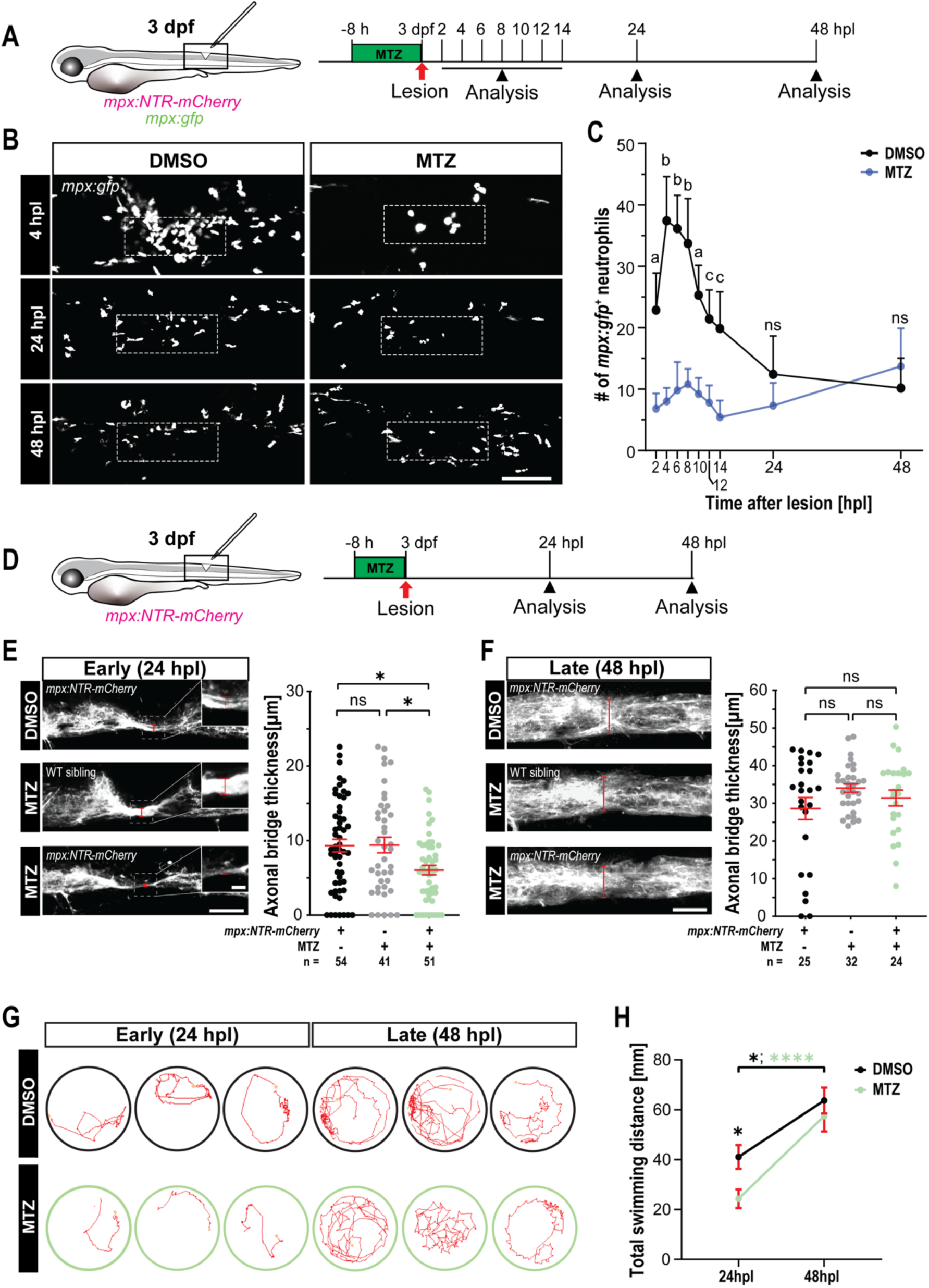
Ablation of neutrophils delays spinal cord repair. **A:** A schematic of a lateral view of a zebrafish larva indicating the spinal cord lesion (boxed) is shown. Rostral is left. Unless otherwise specified, this view is adopted throughout the article. The experimental timeline of nitroreductase/metronidazole mediated neutrophil ablation in *mpx:NTR-mCherry; mpx:gfp larvae*, corresponding to (B,C), is also shown. **B, C:** MTZ treatment significantly reduces neutrophil numbers from 2 to 14 hpl compared to DMSO controls. Dashed boxes mark the quantification area that is centered on the lesion. (Two-way ANOVA, interaction (*F* (8,108) = 14.54, *p* < 0.0001), time (*F* (8,108) = 12.45, *p* < 0.0001), and treatment (*F* (1,108) = 242.9, *p* < 0.0001). Tukey’s multiple comparisons test (DMSO vs. MTZ): a = *p* < 0.001, b = *p* < 0.0001, c = *p* < 0.01, ns indicates no significance. n _DMSO_ = 7, n _MTZ_ = 5). **D:** A schematic timeline for (E-H) is shown. **E, F:** Larvae that underwent neutrophil-ablation (*mpx:NTR-mCherry*+, MTZ+) show reduced axonal bridge thickness, as revealed by anti-acetylated tubulin labelling, at 24 hpl (E) but not at 48 hpl (F), compared to the indicated different control conditions. Dashed boxes mark the areas enlarged in the magnified views of the bridge. Red lines indicate the thickness of the regenerating spinal cord bridge measured for quantification (E: Kruskal-Wallis test, *p* = 0.0126. Dunn’s multiple comparisons test: DMSO (*mpx:NTR-mCherry*) vs. MTZ (*mpx:NTR-mCherry*), \**p* = 0.0229; MTZ (WT sibling) vs. MTZ (*mpx:NTR-mCherry*), \**p* = 0.0491. F: Kruskal-Wallis test, *p* = 0.7796; ns indicates no significance). **G, H:** Larvae that underwent neutrophil-ablation (MTZ) show reduced locomotor activity at 24 hpl, which recovered at 48 hpl, compared to control larvae (H, DMSO). Representative swimming behavior tracks of larvae at 24 and 48 hpl are shown (G). (Two-way ANOVA, interaction (*F* (1,188) = 1.507, *p* = 0.2211), time (*F* (1,188) = 29.08, *p* < 0.0001), and treatment (*F* (1,188) = 6.052, *p* = 0.0148). Tukey’s multiple comparisons test: DMSO (24 hpl vs. 48 hpl), \**p* = 0.0189; MTZ (24 hpl vs. 48 hpl), \*\*\*\**p* < 0.0001; 24 hpl (DMSO vs. MTZ), \**p* = 0.0480). Error bars show SEM. Scale bars: 100 μm (B), 50 μm (E, F), 10 μm (zoom-view of E).

To examine the consequences of neutrophil ablation for spinal cord regeneration, we analyzed axon regrowth, regenerative neurogenesis, and behavioral recovery. Only animals with complete transection of the spinal cord and the notochord left intact entered the experiment. Axon regrowth that reconnects the two spinal stumps was assessed by measuring the thickness of the axonal bundle in the injury site, as previously established (Wehner *et al*, 2017). This indicated a marked reduction in bridge thickness in neutrophil-ablated animals, compared to control groups at 24 hpl (35.2% reduction compared to transgenic animals without MTZ incubation; 35.8% reduction compared to wild type animals with MTZ incubation – controls were not different from each other). At 48 hpl, all groups showed comparable thickness of the bridge, indicating delayed axon regrowth in the absence of the neutrophil peak (Fig. 1D-F).

Similarly, regenerative neurogenesis, measured by EdU incorporation in motor neurons in *mnx1:gfp* reporter animals, as previously established (Cavone *et al*, 2021; Ohnmacht *et al*, 2016), indicated reduced numbers of newly generated motor neurons at 24 hpl compared to the above control groups (Suppl. Fig. 2A-C). At 48 hpl no difference was observed (Suppl. Fig. 2D-E). Ongoing developmental neurogenesis in unlesioned animals was not affected at the corresponding time points, in line with absence of neutrophils during developmental neurogenesis.

Next, we measured the total distance swum after a vibration stimulus of animals at 24 and 48 hpl (Oprişoreanu *et al*., 2023). The distance swum at 24 hpl was 40.8% reduced in MTZ treated *mpx:NTR-mCherry* animals compared to DMSO-treated controls. At 48 hpl, there was no difference anymore, consistent with the dynamics of regeneration and neurogenesis (Fig. 1G,H). Hence, in the absence of the neutrophil peak, axon regrowth, regenerative neurogenesis, and behavioral recovery were delayed, indicating an important role of neutrophils in promoting the regeneration process.

To validate these results with an alternative approach, we blocked neutrophil infiltration of the lesion by inhibiting NADPH oxidase with diphenyleneiodonium chloride (DPI) in an 8 hour drug treatment from 4 hours before to 4 hours after lesion (Niethammer *et al*, 2009). This treatment successfully reduced neutrophil infiltration by 37.9% at 4 hpl. At 24 hpl, 20 hours after wash-out, there was no difference in neutrophil number between control and DPI-treated larvae (Suppl. Fig. 3A-C). The number of Macrophages (labelled by the *mpeg:mCherry* transgene) remained unaffected at 4 and 24 hpl (Suppl. Fig. 3D,E). DPI treatment reduced the thickness of the axon bridge by 49.5% at 24 hpl and by 27.4% at 48 hpl. The reduction of axon regrowth at 24 hpl is in line with the effect of neutrophil ablation. However, the persistent effect at 48 hpl suggests additional effects of the drug on other cell types (Suppl. Fig. 3F-H).

### Ablation of neutrophils changes the inflammatory environment of the lesion site

We reasoned that the lack of neutrophils would affect the cytokine milieu in the lesion site. Therefore, we analyzed changes in immune system regulation by determining levels of expression of pro- (*il1b, tnfa*) and anti-inflammatory cytokines (*tgfb1a, tgfb3*) by qRT-PCR of lesion site-enriched trunk tissue (relative to unlesioned controls) at the peak of neutrophil invasion at 4 hpl, at 24 hpl, when regeneration phenotypes were observed after neutrophil ablation and at the end of our observation period at 48 hpl. We found a persistent approximately 50-80% increase in pro-inflammatory cytokine expression relative to unlesioned controls, but no persistent changes in the anti-inflammatory cytokines tested (Fig. 2A-D). Numbers of Mfap4 immuno-positive macrophages in the injury site remained unchanged in MTZ-treated animals, compared to DMSO-treated controls at 4, 24 and 48 hpl (Fig. 2E-G). This indicated that macrophages could accumulate in the injury site without major neutrophil invasion and that macrophages and potentially other cell types in the lesion site environment showed increased expression of pro-inflammatory cytokines, when neutrophils were ablated (Oprişoreanu *et al*., 2023).

**Figure 2.**
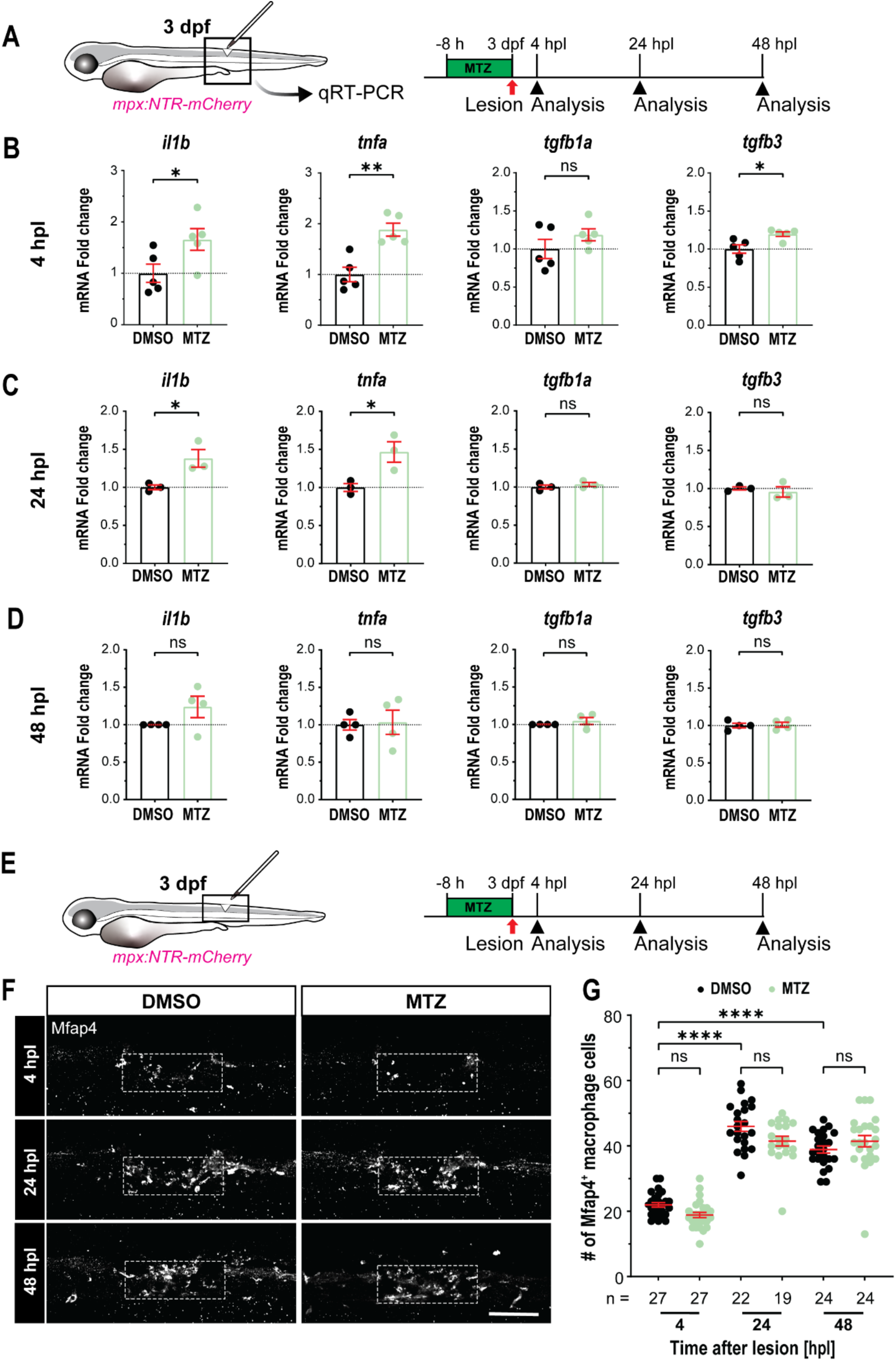
Neutrophil ablation leads to a transient increase in expression of pro-inflammatory cytokines in the lesion site. **A:** A schematic timeline of assessing pro-inflammatory (*il1b, tnfa*) and anti-inflammatory (*tgfb1a, tgfb3*) cytokines expression in *mpx:NTR-mCherry* larvae by qRT-PCR, corresponding to (B-D), is shown. **B, C, D:** Neutrophil ablation leads to a strong upregulation of *il1b* and *tnfa* at 4 and 24 hpl, but not at 48 hpl, with only a slight increase in *tgfb3* at 4 hpl (B). Each dot represents one biological replicate comprising 35 larvae. (Two-tailed unpaired t-test: at 4 hpl, *il1b* (*\*p* = 0.0450), *tnfa* (*\*\*p* = 0.0018), *tgfb3* (*\*p* = 0.0202); at 24 hpl, *il1b* (*\*p* = 0.0338), *tnfa* (*\*p* = 0.0313); ns indicates no significance). **E:** A schematic timeline for (F, G) is shown. **F, G:** Neutrophil ablated larvae (MTZ) show a slight increase in the number of Mfap4⁺ macrophages at 4 hpl, with no significant differences at later time points compared to DMSO controls. Dashed boxes mark the quantification area that is centered on the lesion. (Two-way ANOVA, interaction (*F* (2,137) = 4.464, *p* = 0.0132), time (*F* (2,137) = 217.1, *p* < 0.0001), and treatment (*F* (1,137) = 2.774, *p* = 0.0981). Tukey’s multiple comparisons test: \*\*\*\**p* < 0.0001, ns indicates no significance). Error bars show SEM. Scale bar: 100 μm (F).

### Increased il1b expression delays axonal regeneration after neutrophil ablation

Increased levels of Il-1β have previously been shown to be detrimental to regeneration (Oprişoreanu *et al*., 2023; Tsarouchas *et al*., 2018) and therefore, we aimed to rescue regeneration after neutrophil ablation by disrupting *il1b*. We used a dose-response approach with the caspase-1 inhibitor YVAD, an established drug to reduce Il-1β-signalling by inhibiting caspase-1 dependent processing of the Il-1β protein in zebrafish larvae (Tsarouchas *et al*., 2018).

Indeed, the highest concentration of YVAD (50 µM) exacerbated the effect of neutrophil ablation, whereas a 10 µM concentration restored the thickness of the axon bridge to levels that were not different from those in untreated lesioned control animals. This indicates that the delayed regeneration in neutrophil-ablated animals could at least in part be explained by increased levels of *il1b* expression (Fig. 3A-C).

**Figure 3.**
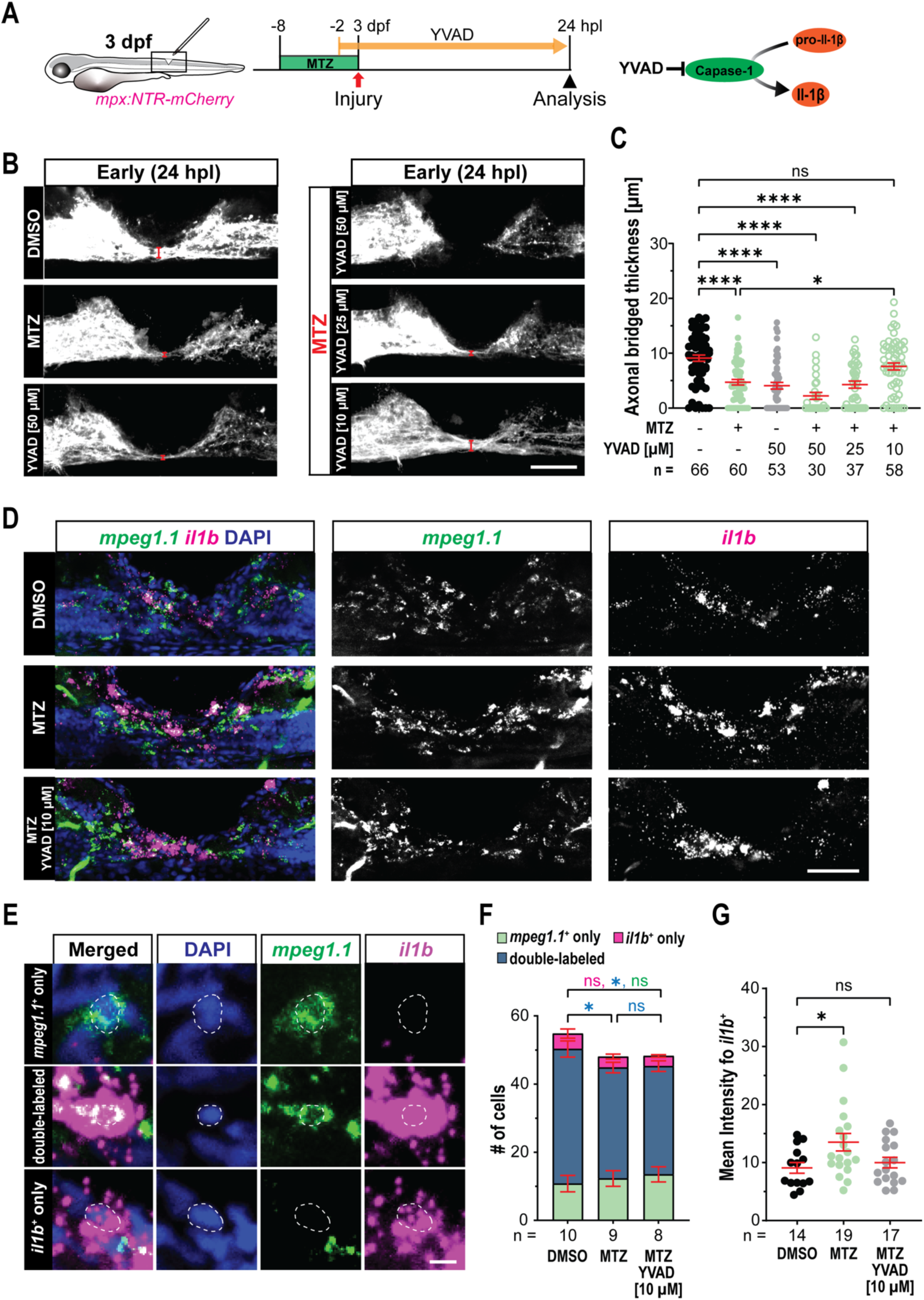
Low-dose inhibition of Il-1β processing restores spinal cord regeneration after neutrophil ablation. **A:** A schematic timeline of YVAD treatment in *mpx:NTR-mCherry* larvae for B-G is shown. **B, C:** 10 µM YVAD treatment restores axonal bridge thickness in larvae that underwent neutrophil ablation, compared to DMSO-treated controls (C). Images of spinal cord lesions labelled with anti-acetylated tubulin in different treatment groups at 24 hpl are shown (B). (Kruskal-Wallis test, *p* < 0.0001. Dunn’s multiple comparisons test: *\*p* = 0.0131, *\*\*\*\*p* < 0.0001, ns indicates no significance). **D, E:** Images of *mpeg1.1* and *il1b* double-HCR in control (DMSO), neutrophil-ablated group (MTZ), and low-dose YVAD-treated neutrophil-ablated (MTZ, 10 µM YVAD) larvae at 24 hpl are shown (D). High-magnification views of individual *mpeg1.1^+^* (green), *il1b^+^*(magenta), and *mpeg1.1^+^*, *il1b^+^* double-labeled cells from D. Dashed lines outline the corresponding cell nucleus **F:** Neutrophil ablation slightly reduces the number of *il1b* expressing macrophages (*mpeg1.1^+^*, *il1b^+^*), compared with the DMSO control. This is not changed by YVAD treatment (Two-way ANOVA, interaction (*F* (4,74) = 2.365, *p* = 0.0606), treatment (*F* (2,74) = 1.517, *p* = 0.2261), and cell category (*F* (2,74) = 223.9, *p* < 0.0001). Tukey’s multiple comparisons test: in *mpeg1.1^+^*, *il1b^+^* double-labeled cells comparisons, DMSO vs. MTZ (*\*p* = 0.0224), DMSO vs. MTZ, 10 µM YVAD (*\*p* = 0.0124), ns indicates no significance). **G:** Neutrophil ablation increases mean *il1b* fluorescence intensity, which is reversed by 10 µM YVAD treatment. (Kruskal-Wallis test, *p* = 0.0530. Dunn’s multiple comparisons test: *\*p* = 0.0402, ns indicates no significance). Error bars show SEM. Scale bar: 50 μm (B, D), 5 μm (E).

### Macrophages/Microglia are the major source of increased il1b expression after neutrophil ablation

To examine the likely cellular source of Il-1β in the lesion core, we counted *il1b*-positive cellular profiles in control, MTZ-treated, and MTZ- and YVAD-treated animals in relation to *mpeg1.1*-expressing macrophages/microglia. We did not consider peripheral *il1b* expression in the skin (Tsarouchas *et al*., 2018). This showed that the vast majority of *il1b*-expressing cells were *mpeg1.1*-positive (between 89.8 and 91.4% of all *il1b+* profiles). About 75% of all macrophages in the injury site expressed *il1b*. There were only minor changes in these ratios between conditions. However, we noticed that the average labeling intensity for *il1b* after MTZ treatment was increased by 60.8% and this was reduced again by co-treatment with 10 µM YVAD to levels that were indistinguishable from the vehicle control (Fig. 3D-G). Reduced *il1b* expression also confirms activity of YVAD, since YVAD treatment has been shown to reduced *il1b* transcription, likely by inhibiting Il-1β positive feedback regulation (Moreno-Loaiza *et al*, 2025; Tsarouchas *et al*., 2018). Hence, ablation of neutrophils leads to increased expression of *il1b*, mainly in macrophages/microglia.

### A subpopulation of neutrophils may signal via Il-4 at the injury site

To determine potential signaling molecules that mediated the pro-regenerative effect of neutrophils on the lesion microenvironment, we interrogated previously generated scRNA-seq data sets on lesioned and unlesioned site tissue at 24 hpl (Docampo-Seara *et al*, 2024). This data set contained all major neural and non-neural tissues, including two clusters of neutrophils (*mpx+*) (Fig. 4A-C). To find selectively expressed signaling molecules in neutrophils, we performed a top-ranked gene expression analysis for these clusters and found high expression levels of *il4* in neutrophil population #2 (Fig. 4D). This expression was highly enriched in that neutrophil cluster compared to all major cell types. Interestingly, most other cell types, including macrophages/microglia, expressed *il4r.1*, a putative receptor for Il-4 (Fig. 4E), making the cytokine a good candidate to mediate neutrophil to macrophage communication.

**Figure 4.**
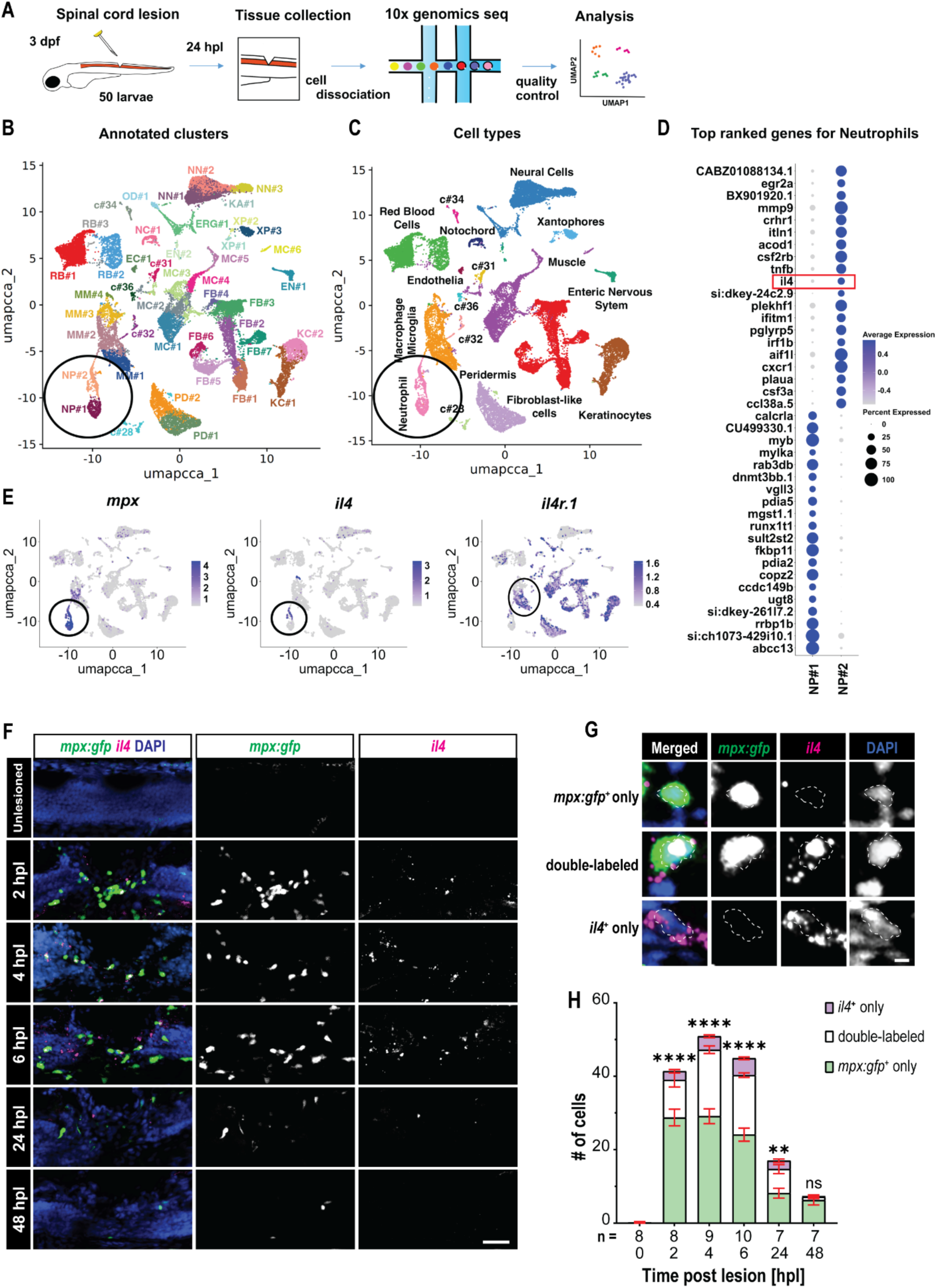
*Il4* is mainly expressed by neutrophils at the spinal cord lesion site. **A:** A schematic of an experimental design for single-cell RNA sequencing (scRNA-seq) is shown. **B, C:** Cluster analysis of scRNA-seq data from the spinal lesion site at 24 hpl is shown (Docampo-Seara *et al*., 2024). Distinct clusters, including two neutrophil clusters (circled), are color-coded and annotated based on marker gene expression. **D:** *il4* ranks among the top 20 most highly expressed genes within the neutrophil sub-population NP#2. **E:** The dot plots identify neutrophils (*mpx;* circled in left and middle panels), show selective expression of *il4* in neutrophils and widespread expression of putative Il-4 receptor *il4r.1* (macrophage/microglia subpopulations are circled in right panel). **F-H:** A time-course analysis of *il4* expression by HCR indicates expression of *il4* mainly in neutrophils (*mpx:gfp^+^*) during spinal cord regeneration (F). High-magnification images show examples of single and double-labelled cells (G). A quantification of *il4* expression is shown in H (Two-way ANOVA, interaction (*F* (10,128) = 24.35, *p* < 0.0001), time (*F* (5,128) = 109.3, *p* < 0.0001), and cell category (*F* (2,128) = 201.1, *p* < 0.0001). Tukey’s multiple comparisons test: in *mpx:gfp^+^*, *il4^+^* double-labelled cells comparisons, 0 h vs. 2, 4 or 6 hpl (*\*\*\*\*p* < 0.0001), 0 h vs. 24 hpl (*\*\*p* = 0.0043), 0 h vs. 48 hpl (ns indicates no significance)). Error bars show SEM. Scale bar: 50 μm (F), 5 μm (G).

To verify expression of *il4* in neutrophils, we performed *in situ* detection of *il4* mRNA in neutrophil reporter animals (*mpx:gfp*) in unlesioned animals at 3 dpf, and after injury at 2, 4, 6, 24, 48 hpl. Quantifications indicated that il4 expression peaked at 4hpl and that a very high proportion of all *il4* expressing cells were neutrophils (*mpx:gfp+*), especially during the peak of neutrophil invasion (At 2 hpl, 81.1 %, at 4 hpl 83.2 %, at 6 hpl 77.9 %, at 24 hpl 74.2 %, and at 48 hpl 75 %). Conversely, at 2 hpl, 26.3 %, at 4 hpl 38.4 %, at 6 hpl 40.2 %, at 24 hpl 44.7 %, and at 48 hpl 12 % of all neutrophils were *il4-positive.* The neutrophils that were negative for an *il4* signal, possibly corresponded to neutrophil population NP#1 seen in scRNA-seq not to express *il4* (Fig. 4F-H).

Ablation of neutrophils also reduced detectability of *il4* by qRT-PCR in unlesioned animals (28.7 %) and showed a tendency to reduction at 4 hpl (21.6 %), but not at 24 and 48 hpl, when neutrophil numbers were back to control levels (Suppl. Fig. 4A,B). The *il4* mRNA HCR analysis following neutrophil ablation also demonstrated an 80.46% reduction in the number of *il4^+^*neutrophils at 6 hpl. At 24 hpl, when overall neutrophil numbers were not different anymore from controls in our previous assays (cf. Fig. 1A,B), there was still a very small reduction in the number of *il4^+^* neutrophils (Suppl. Fig. 4C-G). Hence, *il4* was enriched in specific neutrophils in the lesion and neutrophil ablation reduced abundance of *il4* expression.

### Loss of il4 delays spinal cord regeneration

To directly test the importance of *il4* for spinal cord regeneration, we generated somatic mutants by injecting highly active CRISPR gRNAs into eggs following a previously published protocol (Keatinge *et al*, 2021). Gene disruption in somatic mutants was very efficient, as indicated by the observation that a targeted restriction enzyme recognition site in the coding sequence became almost completely resistant to digestion, as shown by restriction fragment length polymorphism analysis (Suppl. Fig. 5A). Moreover, mRNA expression was also strongly reduced, as indicated by qRT-PCR analysis (up to 75%; Suppl. Fig. 5B). To further validate results, we used a previously published *il4* germline mutants (Bottiglione *et al*, 2020).

Numbers of neutrophils and macrophages in the CHT, expression of major cytokines (*il1b, tnfa, tgfb1a, tgfb3*) and overall morphology were not affected in unlesioned *il4* somatic mutants (Suppl. Fig. 5C-J), indicating that lack of *il4* function did not lead to major aberrations of morphological development and development of the immune system.

To assess effectiveness of axonal regeneration in somatic *il4* mutants, we measured axonal bridge thickness as before. This revealed a 38.1% reduction at 24 hpl compared to control gRNA-injected larvae, with no significant difference observed anymore at 48 hpl (Suppl. Fig.6A-D). Similarly, homozygous *il4* germline mutants showed a 26.6% reduced thickness of the axon bridge at 24 hpl, but no difference anymore at 48 hpl. Heterozygous siblings were unaffected (Fig. 5A-C). This indicated a delay in axon regrowth when *il4* was disrupted, similar to the effect of neutrophil ablation.

**Figure 5.**
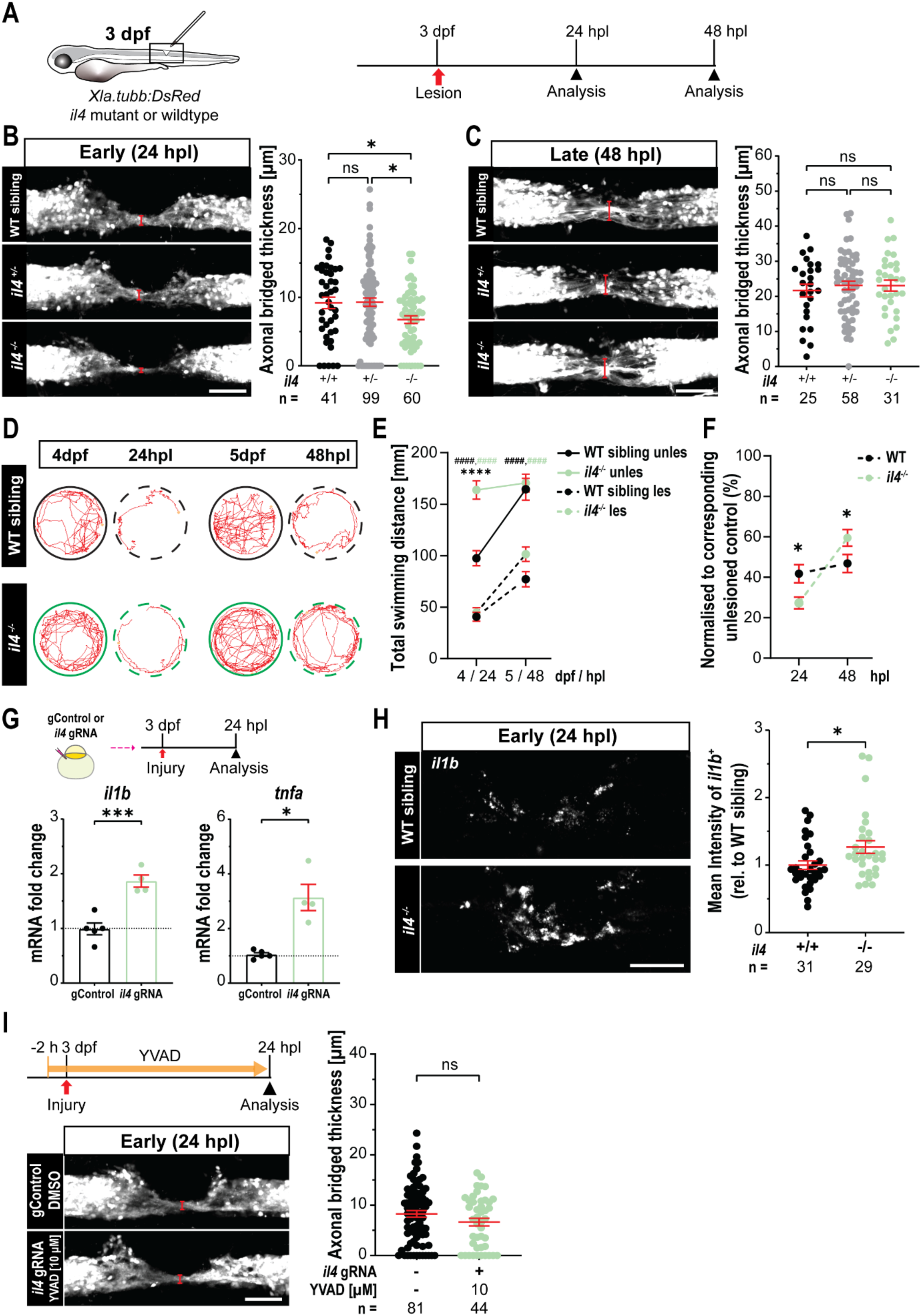
*Il4* disruption delays spinal repair by increasing expression of *il1b*. **A:** A schematic timeline for assessing spinal cord regeneration in *il4* mutants with transgenically labelled neurons (*Xla.tubb:DsRed* fish) corresponding to (B-F, and I), is shown. **B, C:** In *il4* homozygous mutants (*il4^-/-^*), axonal bridge thickness is reduced, compared to wild type (WT) or heterozygote (*il4^+/-^*) siblings at 24 hpl (B), but not at 48 hpl (C). (B: Kruskal-Walls test, *p* = 0.01. Dunn’s multiple comparisons test: *il4^+/+^* vs. *il4^-/-^*(*\*p* = 0.0461), *il4^+/-^* vs. *il4^-/-^* (*\*p* = 0.0154), ns indicates no significance. C: One-way ANOVA, *p* = 0.7822. Tukey’s multiple comparisons test: ns indicates no significance). **D, E:** Unlesioned *il4^-/-^*larvae exhibit greater locomotor activity compared to unlesioned WT siblings at 4 dpf. The lesion causes a significant reduction in swimming distance compared to unlesioned larvae (E). Representative swimming behavior tracks of WT and *il4^-/-^* sibling larvae at different time points are shown (D). (n WT sibling unlesioned = 28, n *il4-/-* unlesioned = 47, n WT sibling lesioned = 42, n *il4-/-* lesioned = 54. Two-way ANOVA, interaction (*F* (5,515) = 4.681, *p* = 0.0003), time (*F* (1,515) = 79.68, *p* < 0.0001), and treatment (*F* (5,515) = 83.58, *p* < 0.0001). Tukey’s multiple comparisons test: unlesioned vs. lesioned (*^####^p* < 0.0001), WT sibling unlesioned vs. *il4-/-* unlesioned (*\*\*\*\*p* < 0.0001)). **F:** Mutant larvae for *il4* show reduced recovery in swimming distance compared to wildtype at 24 hpl (normalized to respective unlesioned control; Two-way ANOVA using a mixed-effects model, interaction (*F* (1,28) = 15.83, *p* = 0.0004), time (*F* (1,53) = 0.0238, *p* = 0.8781), and genotype (*F* (1,53) = 30.29, *p* < 0.0001). Šídák’s multiple comparisons test: *\*p* = 0.0148 at 24 hpl, *\*p* = 0.0270 at 48 hpl). **G:** A qRT-PCR analysis shows that *il1b* and *tnfa* mRNA expression are significantly upregulated in *il4* somatic mutants, compared to gControl-injected larvae. Each dot represents one biological replicate in each independent experiment, comprising 35 larvae. (Two-tailed unpaired t-test: *il1b* (*\*\*\*p* = 0.00090), *tnfa* (*\*p* = 0.0210)). **H:** The *il4^-/-^* larvae increase the mean intensity of *il1b* fluorescence (HCR) at 24 hpl. (Two-tailed unpaired Mann–Whitney test: *\*p* < 0.0194). **I:** 10 µM YVAD treatment rescues axonal bridge thickness in *il4* somatic mutant compared to wild type at 24 hpl. (Two-tailed unpaired Mann–Whitney test: ns indicates no significance). Error bars show SEM. Scale bar: 50 μm (B, C, H, I).

Next, we determined recovery of swimming function in *il4* mutants. We could not find differences to wildtype siblings at 24 and 48 hpl. However, we noticed that unlesioned *il4* mutants exhibited ∼70% longer swim distances than their wildtype siblings, indicating a possible effect of Il-4 on motor system development. Studies in mammals have shown synaptic alterations and hyper-excitability following *Il-4* disruption, which may be related to our observation (Chen *et al*, 2020; Guedes *et al*, 2023) (Fig. 5D,E). Therefore, we normalized swim performance of lesioned larvae to that of their respective unlesioned siblings. We found a relatively worse recovery of *il4* mutants at 24 hpl, compared to wildtype animals. At 48 hpl, their relative swimming capacity was even better than that of wildtype animals (Fig. 5F). This data suggested reduced recovery of axonal bridging and swimming behaviour in *il4* mutant animals at 24 hpl, similar to the effects of neutrophil ablation.

### Rescue of regeneration by reducing increased il1b levels in il4 mutants

To determine possible causes for the delayed regeneration when *il4* is disrupted, we characterized the immune response. First, we analyzed numbers of neutrophils and macrophages after spinal cord injury at 4, 24, and 48 hpl. We found no changes at early timepoints, but increased presence of neutrophils (78.6%) and macrophages (52.8%), at 48 hpl (Suppl. Fig. 6E-H) in somatic *il4* mutants, indicating impaired resolution of inflammation after spinal cord injury when *il4* was disrupted.

To find out whether lack of *il4* expression would increase the abundance of pro-inflammatory cytokines in the lesion, similar to what was observed after neutrophil ablation, we determined expression levels of *il1b* and *tnfa* in *il4* somatic mutants. Indeed, qRT-PCR indicated a doubling of *il1b* expression and three-fold more *tnfa* expression (Fig. 5G). In addition, using *in situ* detection of *il1b* mRNA, we found a 24.2% increase in labelling intensity in *il4* germline mutant animals compared wildtype siblings at 24 hpl (Fig. 5H).

Next, we tested whether inhibiting the elevated Il-1β release with YVAD could rescue axon regrowth in *il4* germline mutants. Indeed, the reduced thickness of the axonal bridge in lesioned mutants was rescued to that observed in wildtype animals by exposing the mutant larvae to 10 µM YVAD, as after neutrophil ablation (Fig. 5I). Hence, changes measured in axon regeneration and immune response in *il4* mutants were similar to those seen after neutrophil ablation and could also be reversed by inhibiting Il-1β processing.

### Il4 over-expression rescues lack of neutrophils

Based on the similarities between the phenotypes of neutrophil ablation and *il4* mutation, we hypothesized that Il-4 signaling may be the major mechanism by which neutrophils promote regeneration. To test this hypothesis, we aimed to rescue delayed regeneration in animals in which neutrophils were ablated by over-expressing *il4*. We did this by using a heat-shock inducible plasmid that was injected into the fertilized egg in a transient transgenesis approach. A single heat-shock at 2 hours before injury led to a ∼100-fold increase in *il4* mRNA detected by qRT-PCR in lesion site tissue at 24 hpl (Suppl. Fig. 7A,B). Using *in situ* detection of *il4* mRNA, we observed that the labelling intensity at 4 and 24 hpl increased by 12-fold and 2.5-fold respectively in animals injected with the *il4* overexpression plasmid, compared to the neutrophil ablation group without injection of the *il4* overexpression plasmid (Suppl. Fig. 7C-F). These observations indicated that after a single heat-shock the plasmid injection led to high levels of *il4* over-expression during the relevant experimental time period.

Next, we measured axonal bridge thickness as a measure of regenerative success in larvae in which we combined neutrophil ablation with *il4* over-expression. Over-expression of *il4* alone after spinal lesion did not change the axonal bridge thickness compared to non-heat shocked and non-MTZ treated controls. Ablation of neutrophils with MTZ treatment alone, reproduced the previously described reduction in axonal bridge thickness at 24 hpl (cf. Fig. 1E). The combination of neutrophil ablation and heat-shock induced over-expression of *il4* increased the thickness of the axonal bridge to values that were not distinguishable anymore from those in controls (Fig. 6A-C). This demonstrated a full phenotypic rescue of reduced axon regrowth caused by neutrophil ablation by over-expression of *il4*.

**Figure 6.**
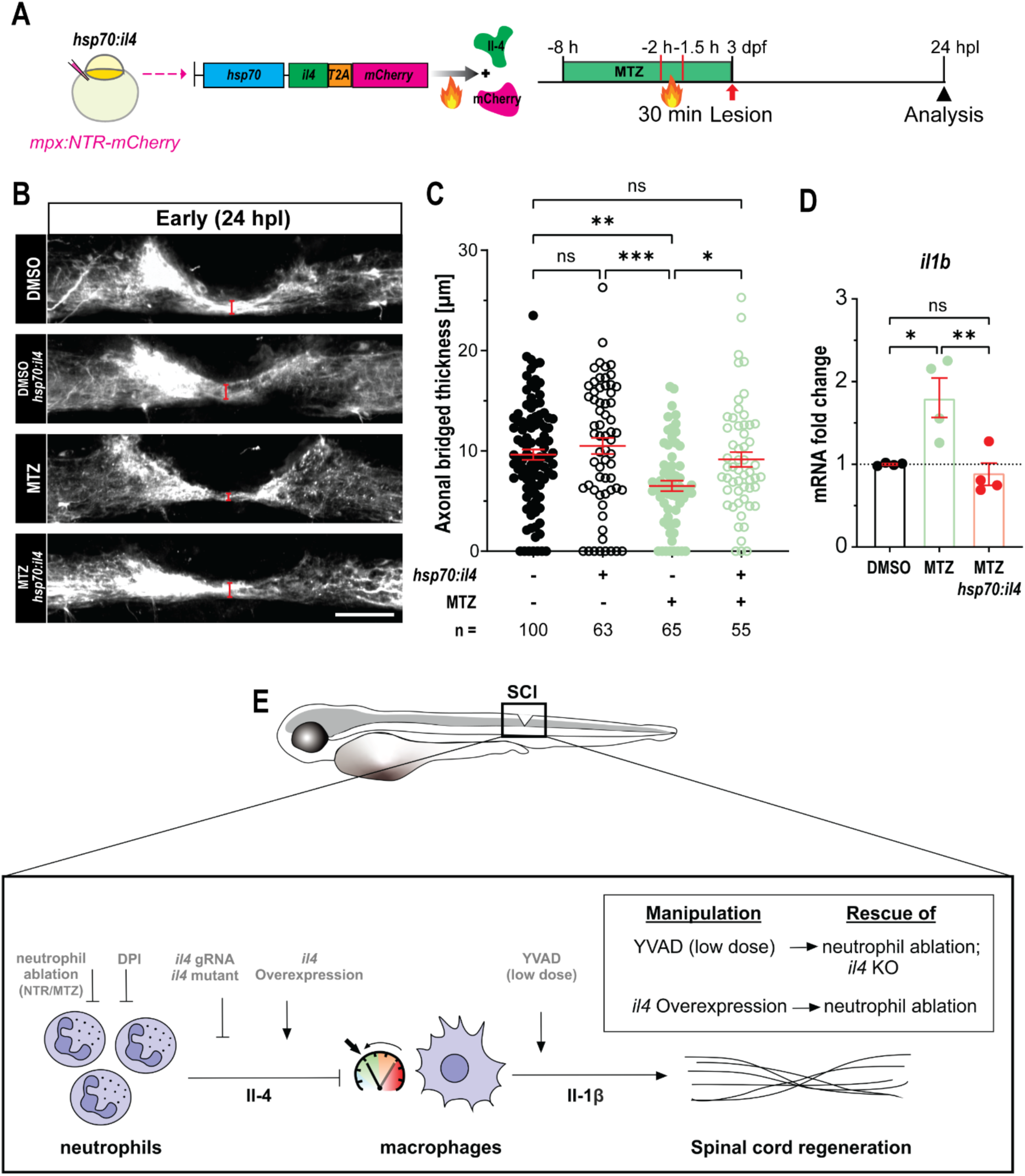
Overexpression of *il4* rescues spinal cord regeneration after neutrophil ablation. **A:** A schematic indicating the *hsp70:il4* plasmid construct and experimental timeline for (B-D) is shown. **B, C:** Heat-shock-induced *il4* overexpression increases the thickness of the regenerative axonal bridge in neutrophil-ablated larvae at 24 hpl, as revealed by anti-acetylated tubulin labelling, compared to neutrophil ablation alone, to that observed in non-heat shocked and non MTZ-treated controls. (Two-way ANOVA, interaction (*F* (1,279) = 1.851, *p* = 0.1748), *hsp70:il4* (*F* (1,279) = 7.316, *p* = 0.0073), and MTZ treatment (*F* (1,279) = 11.88, *p* = 0.0007). Tukey’s multiple comparisons test: *\*p* = 0.0363, *\*\*p* = 0.0016, *\*\*\*p* = 0.0002, ns indicates no significance). **D:** A qRT-PCR analysis shows that heat-shock-induced *il4* overexpression reduces *il1b* expression in neutrophil-ablated larvae to those in control larvae. (One-way ANOVA, *p* = 0.0050. Tukey’s multiple comparisons test: DMSO vs. MTZ (*\*p* = 0.0143), DMSO vs. MTZ *hsp70:il4* (*\*\*p* = 0.0064), ns indicates no significance). **E:** As schematic of the proposed promotion of regeneration by neutrophils via control of macrophage *il1b* expression is shown. Experimental manipulations are depicted in grey and the table indicates rescue experiments. Bars show SEM. Scale bar: 50 μm (B).

To determine whether *il4* overexpression would also rescue the increased expression of pro-inflammatory cytokines after neutrophil ablation, we performed qRT-PCR measurements of *il1b* mRNA in the lesion. Neutrophil ablation led to a ∼2-fold increase in expression levels of *il1b*, compared to vehicle-treated controls at 24 hpl, as before (cf. Fig. 2A,B). Over-expression of *il4* in neutrophil-ablated larvae, reduced the expression level of *il1b* to that seen in vehicle treated, non-heat-shocked controls (Fig. 6D). Hence major changes in regenerative axon growth and in the cytokine milieu were rescued by over-expression of *il4* alone. This shows that neutrophils accelerate regeneration mainly by *il4* expression which reduces *il1b* expression in macrophages (Fig. 6E).

## DISCUSSION

Here we identify a pro-regenerative sub-population of neutrophils in zebrafish. Using a series of rescue experiments, we propose that these neutrophils promote axon regrowth and recovery of swimming behavior mainly via Il-4 signaling, which in turn controls *il1b* expression in macrophages.

Our data suggest that neutrophil-derived Il-4 induces a pro-regenerative phenotype in macrophages. In mouse spinal cord injury, it has been shown that initial levels of IL-4 after spinal cord injury are very low, suggesting that an *il4*-expressing subpopulation of neutrophils is very small, if it exists. Importantly, exogenous supply of IL-4 leads to improved functional recovery, reduced tissue damage and increased expression of tissue repair markers by macrophages (Fenn *et al*, 2014; Francos-Quijorna *et al*, 2016). Similar observations have been made in a myocardial infection model, where IL-4 stimulated anti-inflammatory phenotypes in both macrophages and neutrophils (Daseke *et al*, 2020). In a zebrafish, in a model of hair cell regeneration in the lateral line nerve, macrophage-derived Il-4 promotes synaptogenesis (Denans *et al*, 2022). Il-4 also promotes neurogenesis in a neurodegeneration model in the adult zebrafish brain (Bhattarai *et al*, 2016). These examples indicate direct and indirect pro-regenerative roles of Il-4.

In mammals, neutrophils rarely secrete IL-4 (Brandt *et al*, 2000; Gideon *et al*, 2019). However, in a mouse model of neonatal encephalopathy, caused by hypoxia-ischemia, a late-arriving subpopulation of neutrophils has recently been discovered that promotes repair and expresses *Il-4*, but the functional role of the cytokine has not been tested in that model (Richter *et al*, 2025).

Il-4 from neutrophils could act on macrophages via the Il4r.1 receptor, which is expressed by macrophages, but also via other regeneration-relevant cell types, such as *il4r.1* expressing fibroblasts (Tsata *et al*, 2021). The potentially complex down-stream actions of Il-4 in zebrafish warrant further investigations.

Neutrophils may also have other positive functions for regeneration. For example, in mammals, some neutrophils secrete growth factors, such as NGF, IGF-1 or Oncomodulin, which promote recovery after optic nerve or spinal cord lesion (Kurimoto *et al*, 2013; Sas *et al*., 2020). Neutrophils can also form extracellular material in a regeneration context that may aid regeneration (Bludau *et al*, 2024). Nevertheless, in the larval zebrafish injury model, exogenous Il-4 was sufficient to replace the presence of neutrophils in the injury site, indicating that Il-4 plays a decisive role in neutrophil signalling.

Il-1β is a major determinant of regenerative success. Perturbations of the immune response after injury, e.g., in macrophage-less *irf8* mutants, lead to increased levels of *il1b* expression, which is detrimental to regenerative success and can be rescued by interfering with Il-1β processing directly, or indirectly by treatment with anti-inflammatory drugs, both of which rescue regeneration (Oprişoreanu *et al*., 2023; Tsarouchas *et al*., 2018). In mammals, IL-1β receptor antagonist treatment leads to reduced inflammation and neutrophil recruitment to a spinal cord injury site (Yates *et al*, 2021). However, levels and timing of Il-1β presence in the injury site are critical for regenerative success. Here we show that neutrophils control expression of *il1b* in macrophages and that reducing Il-1β processing to the right dose is critical for rescue experiments. Previously, we observed that reducing Il-1β processing during early, but not late phases of regeneration in wildtype animals impaired axon regrowth, such that early expression of *il1b*, likely from neutrophils themselves, promotes regeneration (Tsarouchas *et al*., 2018).

Lack of neutrophils also leads to increased expression of *tnfa*, which promotes axon regrowth and regenerative neurogenesis on its own (Cavone *et al*., 2021; Tsarouchas *et al*., 2018). Apparently, the positive functions of *tnfa* are outweighed by the negative ones of *il1b* for regeneration. There are also complex feedback interactions between these two cytokines, as *tnfa* is under positive control by *il1b* and *il1b* is under negative control by *tnfa* in the lesion site (Tsarouchas *et al*., 2018).

A potential technical limitation of our study is that macrophages, which are highly phagocytotic after spinal cord injury, also phagocytose nitroreductase-ablated neutrophils, leading to a higher proportion of neutrophil-loaded macrophages than in controls. Neutrophil phagocytosis by macrophages can lead to altered reactions to IL-4 (Liebold *et al*, 2024). This potential confounder is mitigated by our observation that preventing neutrophil infiltration with DPI led to similar regeneration phenotypes.

Moreover, when only *il4* is disrupted without killing the neutrophils, the resulting regeneration and cytokine milieu changes are comparable to those observed after neutrophil ablation. In particular, in both situations *il1b* expression is increased and phenotypes are fully rescued by reducing Il-1β processing.

We propose that there are conserved aspects in the innate immune response to spinal cord injury between larval zebrafish and mammals in that IL-4 reprograms macrophages to a pro-repair phenotype. A notable difference is that IL-4-expressing neutrophils are rare in the initial immune response in mammals, but not in zebrafish.

Therefore, this novel sub-population of *il4-*expressing neutrophils in larval zebrafish could partially explain the remarkable regenerative success in that species. Our findings also support the notion that therapeutic efforts in mammals may be directed to inducing more pro-regenerative IL-4-producing neutrophils.

## MATERIALS AND METHODS

### Animals

All zebrafish lines were raised and kept under standard conditions (Brand *et al*, 2002; Westerfield, 2007). Zebrafish experiments were performed under state of Saxony licenses TVV 36/2021, TVV 45/2018, and holding licenses DD24-5131/364/11, DD24-5131/364/12. For experimental conditions, larvae up to an age of 5 days post-fertilization (dpf) were used of the following lines: wild type (AB); *Tg(mpx:GFP)^i114^*, abbreviated as *mpx:gfp* (Renshaw *et al*, 2006); *TgBAC(mpx:GAL4-VP16)^sh267^*, *Tg(UAS-E1B:NTR-mCherry)^c264^*, abbreviated as *mpx:NTR-mCherry* (Davison *et al*, 2007; Robertson *et al*, 2014); *Tg(mnx1:GFP)^ml2^*, abbreviated as *mnx1:gfp* (Flanagan-Steet *et al*, 2005); *Tg(mpeg1.1:mCherry)^gl23^*, abbreviated as *mpeg:mCherry* (Ellett *et al*, 2011); *Tg(Xla.Tubb:DsRed)^zf148^*, abbreviated as *Xla.Tubb:DsRed* (Peri & Nusslein-Volhard, 2008); *il4^umc18/umc18^*, abbreviated as *il4* mutants (Bottiglione *et al*., 2020). For all experiments, zebrafish larvae were kept in E3 medium (Brand *et al*., 2002) without methylene blue supplement.

### Image acquisition, processing, and presentation

Images were acquired using the systems described in each subsection. Images were processed using ImageJ (http://rsb.info.nih.gov/ij/; version 2.3.0/1.54), and Adobe Photoshop 2025 (version 25.6), and Figure plates were prepared using Adobe Illustrator 2024 (version 28.7.5). Images of the whole animal, zebrafish trunk or lesion site are lateral views (dorsal up; rostral left), if not indicated differently.

### Spinal cord lesion

Lesions of larvae were performed as previously described (Ohnmacht *et al*., 2016). Briefly, at 3 dpf, zebrafish larvae were anaesthetized in E3 medium (Nüsslein-Volhard, 2002) containing 0,02 % of MS-222 (Sigma-Aldrich, St. Louis, MO, USA; Cat#A5040). Larvae were then transferred to an agarose plate. After removal of excess water, larvae were placed in lateral positions. A transection of the entire spinal cord was made using a 30G x ½″ hypodermic needle (Becton Dickinson, Franklin Lakes, NJ, USA; Cat#613-3942) at the level of the urogenital pore, without injuring the notochord. After surgery, larvae were returned to E3 medium for recovery and kept at 28.5°C. Larvae in which the notochord had been inadvertently damaged were excluded from further analysis.

### CRISPR/Cas9 manipulation

CRISPR/Cas9-mediated mutagenesis was achieved following a previously published protocol (Keatinge *et al*, 2021). CRISPR gRNAs were selected using the CRISPRscan (www.crisprscan.org/gene) web tool (Moreno-Mateos *et al*, 2015), and the CRISPOR (http://crispor.org) web tool (Concordet & Haeussler, 2018). gRNAs were chosen based on the CRISPRScan score (preferentially > 60) and the CRISPOR scores (high specificity, high efficiency and low out-of-frame scores), and on the availability of a suitable restriction enzyme recognition site (www.snapgene.com) that would be destroyed by the gRNA. Additional criteria were to avoid the first exon and to target functional domains. All gRNAs and tracrRNA were ordered from Merck KGaA (Darmstadt, Germany). 1 nL of gRNA mix (1 µL of Tracer (5nM, Merck KGaA, Darmstadt, Germany; Cat#TRACRRNA05N), 1 µL of each gRNA (20µM,Merck KGaA, Darmstadt, Germany), 1 µL of phenol red (Sigma-Aldrich, St. Louis, MO, USA; Cat#P0290) and 1 µL of Cas9 (20µM, New England Biolabs, Ipswich, MA, USA; Cat#M0386M) was injected into one-cell stage embryos. gRNA efficiency was tested by restriction fragment length polymorphism analysis (RFLP) in a proportion of larvae for each experiment as an internal experimental control (Keatinge *et al*., 2021). Sequences for gRNAs, primers and the restriction enzymes (RE, New England Biolabs, Ipswich, MA, USA) used for RFLPs are provided in Supplementary Table 1. The *il4* gene (ENSDARG00000087909) was targeted by injection of gRNA for exon 4 (5’-CTTCATTGTGCATTCCCCCG -3’). All experiments conducted with somatic mutants were carried out with a control injected group with a non-targeting gRNA (5’-UUACCUCAGUUACAAUUUAU -3’).

### Generation of heat shock plasmid and treatment

To create the *pTol-hsp70l:il4-T2a-mCherry* plasmid, we modified the *pTol-hsp70I:sema4ab-T2a:mCherry* vector (Docampo-Seara *et al*, 2025). We PCR-amplified the *il4* coding sequence from genomic cDNA with specific primers (Supplementary Table 2). Then, we performed a simultaneous multiple fragment ligation in the opened backbone vector. The final construct *pTol-hsp70l:il4-T2a:mCherry* was verified by sequencing, qRT-PCR for over-expression and *il4* mRNA fluorescence (HCR-FISH) after injection.

### In situ hybridization chain reaction (HCR-FISH)

Hybridization chain reaction (HCR) RNA fluorescence *in situ* hybridization (Molecular Instruments, Los Angeles, CA, USA) was performed according to the manufacturer’s instructions with minor modifications (Choi *et al*, 2018; Schwarzkopf *et al*, 2021). In brief, terminally anesthetized larvae were fixed overnight in 4 % paraformaldehyde (PFA, Thermo Scientific, Waltham, MA, USA; Cat#28908) in phosphate buffered saline (PBS, Mediakitchen of TUD, Dresden, Germany) at 4 °C and then transferred to 100 % Methanol (MeOH, Carl Roth, Karlsruhe, Germany; Cat#4627.1) at −20 °C overnight for permeabilization. The next day, larvae were rehydrated in a graded series of MeOH at room temperature (RT) and subsequent washes in phosphate buffered saline with 0.1 % Tween (PBST). The larvae were treated with 100% acetone (pre-chilled to −20°C. Carl Roth, Karlsruhe, Germany; Cat#7328.1) at −20°C for 10 minutes, and washed with PBST (3 × 5 min). Thereafter, larvae were treated with 1ml proteinase K (Roche Diagnostics GmbH, Mannheim, Germany; Cat#3115828001) at a concentration of 12.5 μg/mL for 60 min at RT, followed by post-fixation in 4 % PFA for 15 min RT and washes with PBST (3 x 5 min). Pre-hybridisation was performed using 500 μL of pre-warmed probe hybridisation buffer (Molecular Instruments, Los Angeles, CA, USA) for 30 min at 37 °C (Choi *et al*., 2018). The probe sets, which were designed based on the NCBI sequence by Molecular Instruments (Choi *et al*., 2018), were prepared using 2 pmol of each probe set in 500 μL of pre-warmed hybridisation buffer. Then, buffer was replaced with probe sets and incubated for 16 h at 37 °C. On the following day, samples were washed 4 x 15 min with pre-warmed probe washing buffer (Molecular Instruments, Los Angeles, CA, USA) at 37 °C followed by 2 x 5 min in 5x sodium chloride sodium citrate with 0.1 % Tween (SSCT) at RT. Larvae were then treated with 500 μL of room temperature equilibrated amplification buffer (Molecular Instruments, Los Angeles, CA, USA) for 30 min at RT. Hairpin RNA preparation was performed following the manufacturer’s instructions.

Samples were incubated with Hairpin RNAs for 16 h in the dark at RT. Finally, samples were washed for 2 x 45 min in 5 x SSCT with DAPI (1:1000, Sigma-Aldrich, St. Louis, MO, USA; Cat#D9564), followed by 3 x 5 min in 5 x SSCT at RT. Samples were mounted in 75 % glycerol (Carl Roth, Karlsruhe, Germany; Cat#3783.1) in PBS and imaged using a Dragonfly spinning disc microscope (Andor/Oxford Instruments, Belfast, UK).

### Combining EdU with motor neuron labelling

Labelling newly generated motor neurons followed an established protocol (Ohnmacht *et al*., 2016). All incubations were performed at room temperature unless stated otherwise. Briefly, lesioned and unlesioned larvae were incubated in E3 medium containing 1 % dimethylsulfoxide (DMSO, Sigma-Aldrich, St. Louis, MO, USA; Cat#D8418) and 1:100 EdU (Click-iT™ Plus EdU Cell Proliferation Kit. Thermo Fisher Scientific, Waltham, MA, USA; Cat#C10640) for the period determined by the experimental design. Terminally anesthetized larvae were fixed in 4 % PFA overnight at 4 °C and permeabilised in 100 % MeOH at −20 °C for at least overnight. After removing the head with micro scissors, larvae were then rehydrated, and incubated in proteinase K and re-fixed in 4 % PFA. Then, larvae were incubated with the Click-iT™ Plus EdU Cell Proliferation Kit (Alexa Fluor™ 647 dye) as described by the manufacturer. Subsequently, samples were washed in PBST and incubated with blocking buffer solution (2 % BSA, 1 % DMSO, 1 % Triton, 10 % Normal donkey serum, in PBST) for 1 h and then with a chicken anti-GFP antibody, (1:200, Abcam, Cambridge, UK; Cat# Ab13970) for 72 hours at 4 °C. Secondary antibody (Alexa Fluor® 488 Donkey Anti-Chicken IgY [IgG], 1:200, Jackson ImmunoResearch Laboratories, West Grove, PA, USA; Cat#703-545-155) incubation took place overnight at 4 °C. Finally, samples were washed in PBST and transferred to 70 % glycerol in PBST, mounted and imaged.

Imaging was performed using a Dragonfly spinning disc microscope. Images were analysed using ImageJ (Rueden *et al*, 2017; Schindelin *et al*, 2012). Newly generated motor neurons were counted by assessing the number of double-positive cells for GFP and EdU. Cells were counted manually along the Z-stack, ensuring counting individual cells in single optical sections, in 50 µm windows located at both sides of the injury site (100 µm in total). To mitigate bias, experimental conditions were occluded for the analysis. At least two independent experiments were performed. Individual experiments were merged and analysed for statistical significance.

### Fluorescent immunohistochemistry

Anti-Mfap4 and anti-Mpx immunohistochemistry on whole-mount zebrafish larvae was carried out as an established protocols (John *et al*, 2022). Briefly, terminally anesthetized larvae were fixed in 4% PFA overnight at 4 °C. After removing the head with micro-scissors, larvae were permeabilized by subsequent incubation in acetone and proteinase K. Samples were re-fixed in 4% PFA, blocked in blocking buffer solution (2 % BSA, 1 % DMSO, 1 % Triton, 10 % Normal donkey serum, in PBST), and incubated over three nights with rabbit polyclonal anti-Mfap4 antibody (GeneTex, Irvine, CA, USA; Cat#GTX132692) or or rabbit polyclonal anti-Mpx antibody (GeneTex, Irvine, CA, USA; Cat# GTX128379) at a dilution of 1:200.

For anti-acetylated tubulin immunolabelling, terminally anesthetized larvae were fixed in 4% PFA for 1 h at room temperature. After removing the head with micro scissors, larvae were permeabilized by subsequent incubation in acetone and proteinase K. Samples were re-fixed in 4% PFA, blocked with blocking buffer solution, and incubated over three nights with mouse monoclonal anti-acetylated tubulin antibody (Sigma-Aldrich, St. Louis, MO, USA; Cat#T6793) at a dilution of 1:300.

The corresponding secondary antibodies conjugated to Alexa Fluor dyes were obtained from Jackson ImmunoResearch (Cat#711-605-152, 715-605-151) and used at a dilution of 1:300 and incubated overnight at 4°C. Finally, samples were mounted with 70 % glycerol in PBST, and imaged using a Dragonfly spinning disc microscope.

### Measurement of axonal regrowth

As an indicator for successful axon regrowth, we determined the extent of continuous labelling of axons across the spinal cord injury site, termed axonal bridging, as previously described (Tsarouchas *et al*., 2018; Wehner *et al*., 2017). We determined the thickness of the axonal bridge at its narrowest point. Spinal lesion was performed at 3 dpf as described in larvae above. Terminally anesthetized larvae were fixed in 4 % PFA and immuno-labelled against acetylated tubulin (if axons were not transgenically labelled). Images were taken with a Dragonfly Spinning Disk microscope with a step size of 1 µm.

The bridge thickness was determined using a custom script in Fiji (Rueden et al., 2017; Schindelin et al., 2012). Measurements were performed at the narrowest point of the axonal bridge, which was manually identified on a maximum intensity projection of the z-stack after binary segmentation. Data obtained from at least three independent experiments were quantified.

### Analysis of swimming behaviour

The swimming behaviour was assessed using the ZebraBox setup (ViewPoint Behavior Technology, Lyon, France). A spinal cord lesion was inflicted at 3 dpf, and its completeness was confirmed at 2 hpl in each larva. Subsequently, the larvae were individually arranged in 24-well plates, ensuring representation of all groups on each plate, and kept in an incubator. Larval movements were automatically tracked.

Measurements encompassed three repetitions of the same protocol. The entire experimental procedure lasted 2 minutes. The first minute served as a rest period to allow the animals to adapt to the device environment. The second minute consisted of a 1-second vibration stimulus (frequency set to 1000 Hz; 100% target power), followed by 59 seconds without stimulation. For analysis, we focused on the full second-minute interval, which included the 1-second stimulus and the subsequent 59-second no-stimulus phase. The analysis was performed with ZebraZoom (https://zebrazoom.org/) (Mirat *et al*, 2013), an open-source tracking software, to track the swimming paths as well as measure the swim distance of each larva. Two independent experiments were performed and the results were merged for statistical analysis.

### Quantitative RT-PCR

RNA was extracted using the Direct-zol RNA Microprep Kit (Zymo Research, Irvine, CA, USA; Cat#R2060) according to the manufacturer’s instructions (40 larvae/condition). The lesioned site or the corresponding unlesioned tissue was enriched by cutting away rostral and caudal parts from the larvae at positions approximately 3 somites away from the lesion site in the rostral and caudal direction. cDNA was synthesized using the Transcriptor High Fidelity cDNA Synthesis Kit (Sigma-Aldrich, St. Louis, MO, USA; Cat#5091284001) according to the manufacturer’s instructions. qRT-PCR was performed using the FastStart Universal SYBR Green Master (Sigma-Aldrich, St. Louis, MO, USA; Cat#4913914001) according to the manufacturer’s instructions and run in triplicates on a LightCycler 480SW instrument (Roche Diagnostics GmbH, Mannheim, Germany). At least three individual experiments were run and the data was analyzed using the LightCycler 96/384 application software v1.1 (Roche Diagnostics GmbH, Mannheim, Germany). Each condition was normalized to housekeeping gene (*β-actin*) and the experimental group was compared to controls by normalizing each value to the control average value. Primers were designed to span an exon–exon junction using Primer-BLAST software or obtained from the literature. Primer sequences are given in Supplementary Table 3.

### Immune cell counting in whole-mounted larvae

Larvae were lesioned at 3 dpf and fixed for 1 h in 4 % PFA at different time points. Larvae were washed in PBST and transferred to glycerol (70 % in PBST), mounted and imaged immediately using a Dragonfly Spinning Disk microscope. Images were analysed using Fiji with the experimental condition unknown to the observers. The number of cells in the vicinity of the injury site was counted in a volume of interested that was centered on the lesion site (width, X-axis - 200 µm, height, Y-axis - 75 µm, depth, Z-axis - 50 µm), right above the notochord. Cells were counted manually inside the window along the Z-stack with the experimental condition occluded. At least two individual experiments were carried out and merged.

### Acridine Orange labelling

Larvae of the *mpx:NTR-mCherry* line were incubated in 1 µg/ml Acridine Orange (Sigma-Aldrich, St. Louis, MO, USA; Cat#A6014) in E3 medium for 3.5 h prior to imaging (Brand *et al*, 1996). Following incubation, fish were washed three times for 10 min each in E3. Larvae were then mounted in 0.5% low-melting-point agarose (Carl Roth, Karlsruhe, Germany; Cat#6351.1) in E3 for live imaging using a Dragonfly spinning disk confocal microscope. To quantify cellular debris, an observation window (width, X-axis = 585 µm; height, Y-axis = 100 µm; depth, Z-axis = 50 µm) was defined over the region of interest centred on the caudal hematopoietic tissue. The number of cells co-expressing mCherry and Acridine Orange signals was counted within this volume.

### Fluorescence intensity measurements

Images were analysed with Fiji ImageJ (https://imagej.net/ij/). The region of interest was defined as a rectangle of 250 x 100 µm, centred on the injury site and located right above the notochord. 50 Z-stacks were collapsed in sum intensity projections and then background subtraction was performed before analysing the mean intensity with a custom Fiji macro to automatize the process (Rueden *et al*., 2017; Schindelin *et al*., 2012). Mean intensity was represented by normalizing each value to the control average value.

### Drug treatment and heat shock

Drug treatments and heat shock were performed according to the schematic timelines shown with the experiment. Larvae were incubated in E3 medium containing the drugs, with a final DMSO concentration of 1 %. To inhibit caspase-1 activation of Il-1β, Ac-YVAD-cmk (Sigma-Aldrich, St. Louis, MO, USA; Cat# SML0429) (Tsarouchas *et al*., 2018) was dissolved in DMSO to a stock concentration of 10mM and used at a final concentration of 10, 25, and 50 µM. To block neutrophil infiltration by inhibiting NADPH oxidase, diphenyleneiodonium chloride (DPI, Sigma-Aldrich, St. Louis, MO, USA; Cat#D2926) (Niethammer *et al*., 2009) was dissolved in DMSO to a stock concentration of 10 mM and used at a final concentration of 100 µM. For targeted cell ablation of *mpx^+^* cells, the Metronidazole (MTZ; MedChemExpress, Monmouth Junction, NJ, USA; Cat#HY-B0318) was dissolved in DMSO and used at a final concentration of 5 mM. For heat shock, larvae were kept in 50-mL conical centrifuge tubes filled with pre-warm E3 medium, which floated in a programmable thermostat-controlled water bath (VWR International BVBA, Leuven, Belgium). Heat shock was performed for 30 min at 38°C, after which larvae were returned to 28.5°C.

### scRNAseq analysis

A previously obtained Seurat object was used (Docampo-Seara *et al*., 2025). Neutrophils were identified by enriched expression of *mpx* and subset for further analysis. For the differential gene analysis the FindAllMarkers function from Seurat (version 5.0.2) was used with default parameters. The “Wilcoxon rank sum test”, and p_val_adj (p-value adjusted) < 0.1 and avg_log2FC >= 0.25 (average log2 fold change) were used for selecting significantly expressed genes. Genes were ordered based on their avg_log2FC in descending order for each neutrophil cluster. The 20 top-ranked genes were then selected based on their representative expression of each cluster vs the other (pct.1 > 0.2, pct.2 < 0.2).

### Bias mitigation and statistical analysis

For all quantifications, investigators were blinded to the experimental conditions. Unless indicated, no data were excluded from analysis. The experiment was independently repeated at least twice, and a two-way ANOVA was used to evaluate treatment effects and to confirm consistency across experiments for data pooling (Montgomery, 2017). All statistical analyses were performed using GraphPad Prism 10 (GraphPad Software Inc., version 10.6.0). All quantitative data were tested for normal distribution using the Shapiro-Wilk test. Parametric and non-parametric tests were used as appropriate. We used two-tailed Welch’s t-test, two-tailed Mann–Whitney test, Kruskal–Wallis test with Tukey’s multiple comparisons test, One-way ANOVA with Tukey’s multiple comparisons test, Two-way ANOVA with Tukey’s multiple comparisons test or Šídák’s multiple comparisons test. Differences were considered statistically significant at p < 0.05. Unless otherwise stated, data are presented as mean ± standard error of the mean (SEM). Graphs were generated using GraphPad Prism 10. Unless otherwise indicated, each data point represents one animal.

## ACKNOWLEDGEMENTS

We thank Drs. Stephen A Renshaw, Didier Stainier and João Cardeira-da-Silva for providing the *TgBAC(mpx:GAL4-VP16)^sh267^*, *Tg(UAS-E1B:NTR-mCherry)^c264^* fish; Drs. Adam Hurlstone and Victoriano Mulero for providing *il4^umc18/umc18^* mutant fish; Drs. Michael Brand and Daniel Wehner for critical reading; and Dr. Judith Konantz, Marika Fischer, Silvio Kunadt, and Denise Bärhold for fish care. Experimental work was supported by the Light Microscopy Facility, the DRESDEN-concept Genome Center, the Flow Cytometry Facility, and the Zebrafish Facility, all core facilities of the Center for Molecular and Cellular Bioengeneering (CMCB) at the Technische Universität (TU) Dresden. Funding was provided by a Chinese Scholarship Council PhD fellowship (to XT, no.202108440240), an Alexander von Humboldt Stiftung Professorship award (to CGB), and TU Dresden core funding (to CGB).

## AUTHOR CONTRIBUTIONS (CRediT nomenclature)

XT: conceptualization, methodology, validation, formal analysis, investigation, writing - original draft, writing - review and editing, visualization; ADS: methodology, validation, formal analysis, investigation, visualization; KH: methodology, validation, formal analysis, investigation, visualization; FK: software; methodology; DZ: investigation; AB: investigation; TB: conceptualization, writing - original draft, writing - review and editing, visualization, supervision, funding acquisition; CGB: conceptualization, writing - original draft, writing - review and editing, visualization, supervision, funding acquisition.

## SUPPLEMENTARY FIGURES

**Suppl. Fig. 1.**
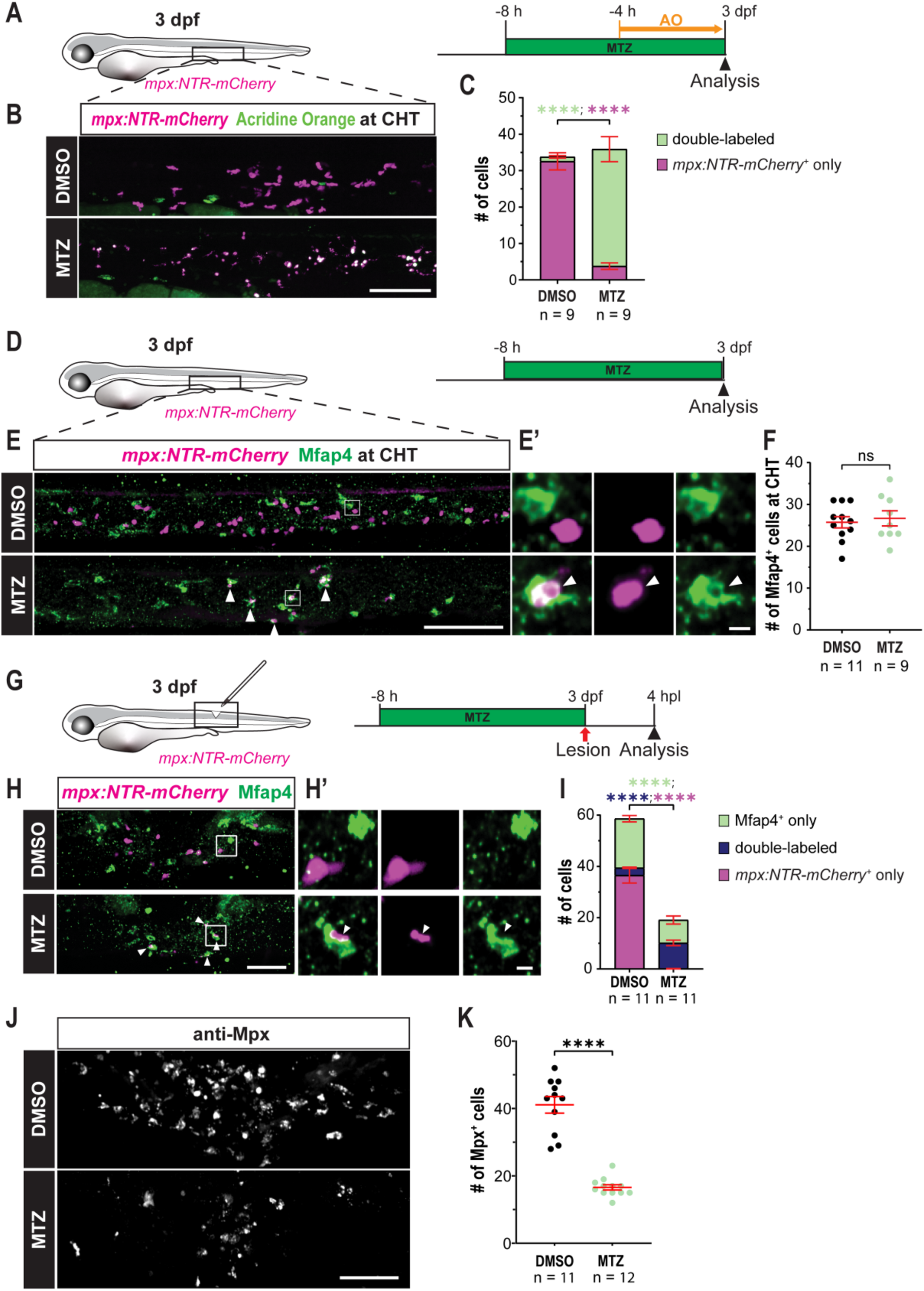
MTZ treatment efficiently ablates neutrophils. **A:** A schematic timeline of MTZ treatment and Acridine Orange (AO) staining for (B, C) is shown. **B, C:** Approximately 89.7% of neutrophils (magenta) are acridine orange positive (green) after MTZ treatment in the caudal hematopoietic tissue (CHT) (Two-way ANOVA, interaction (*F* (1,32) = 195.0, *p* < 0.0001), treatment (*F* (1,32) = 0.2442, *p* = 0.6246), and cell category (*F* (1,32) = 0.4931, *p* = 0.4876). Tukey’s multiple comparisons test: *\*\*\*\*p* < 0.0001). **D:** A schematic timeline of anti-Mfap4 staining in unlesioned larvae for (E-F) is shown. **E, E’:** The *mpx:NTR-mCherry⁺* neutrophils (magenta) and Mfap4⁺ macrophages (green) in the CHT of DMSO- and MTZ-treated larvae. Arrowheads mark Mfap4⁺ macrophages engulfing neutrophil debris. **F:** The numbers of Mfap4⁺ macrophages are not significantly different between DMSO- and MTZ-treated groups in the CHT area. (Two-tailed unpaired t-test, ns indicates no significance). **G:** A schematic timeline of anti-Mfap4 staining in lesioned larvae for (H-I) is shown. **H, I:** MTZ treatment increases the number of *mpx:NTR-mCherry⁺*/ Mfap4⁺ cells at 4 hpl in the lesioned area. (Two-way ANOVA using a mixed-effects model, interaction (*F* (1.827,36.54) = 102.7, *p* < 0.0001), treatment (*F* (1,20) = 102.9, *p* < 0.0001), and cell category (*F* (1.827,36.54) = 30.51, *p* < 0.0001). Tukey’s multiple comparisons test: *\*\*\*\*p* < 0.0001). **J, K:** MTZ treatment of *mpx:NTR-mCherry* transgenic animsl significantly reduced the number of anti-Mpx^+^ neutrophils at 4 hpl in the lesioned area. (Two-tailed unpaired t-test: *\*\*\*\*p* < 0.0001). Error bars show SEM. Scale bars, 100 μm (B, E), 50 μm (H,J) and 5 μm (E’, H’).

**Suppl. Fig. 2.**
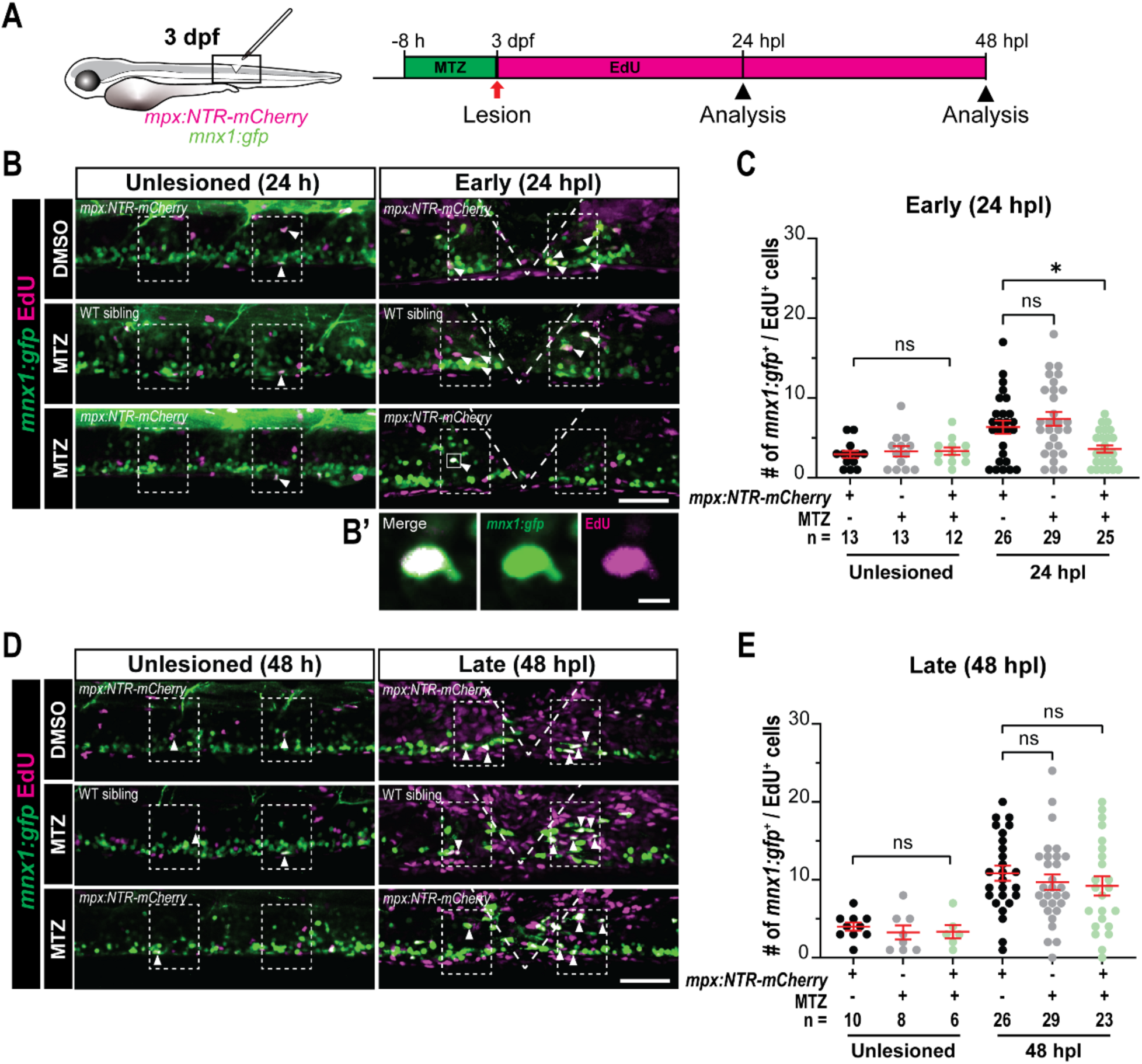
Neutrophil ablation delays regenerative neurogenesis. **A:** A schematic timeline of assessing motor neuron proliferation for (B-E) is shown. **B, C:** Neutrophil ablation reduces the number of newly generated motor neurons (*mnx1:gfp⁺*/ EdU⁺) at 24 hpl, compared to the indicated different control conditions. MTZ treatment doesn’t affect the developmental neurogenesis of motor neuron at 4 dpf. Dashed squares indicate the region of quantification; arrowheads indicate double-labeled cells. High-magnification images of a double-labeled neuron (*mnx1:gfp⁺*/ EdU⁺) (B’), (Two-way ANOVA, interaction (*F* (2,112) = 3.066, *p* = 0.0505)), treatment (*F* (2,112) = 2.686, *p* = 0.0725)), and time (*F* (1,112) = 15.21, *p* = 0.0002)). Tukey’s multiple comparisons test: *\*p* = 0.0480, ns indicates no significance). **D, E:** MTZ treatment does not affect the motor neuron neurogenesis (*mnx1:gfp⁺*/ EdU⁺) at 5 dpf and 48 hpl. (Two-way ANOVA, interaction (*F* (2,96) = 0.05609, *p* = 0.9455), treatment (*F* (2,96) = 0.4050, *p* = 0.6681), and time (*F* (1,96) = 30.53, *p* < 0.0001). Tukey’s multiple comparisons test: ns indicates no significance). Error bars show SEM. Scale bars, 50 μm (B, D) and 5 μm (B′).

**Suppl. Fig. 3.**
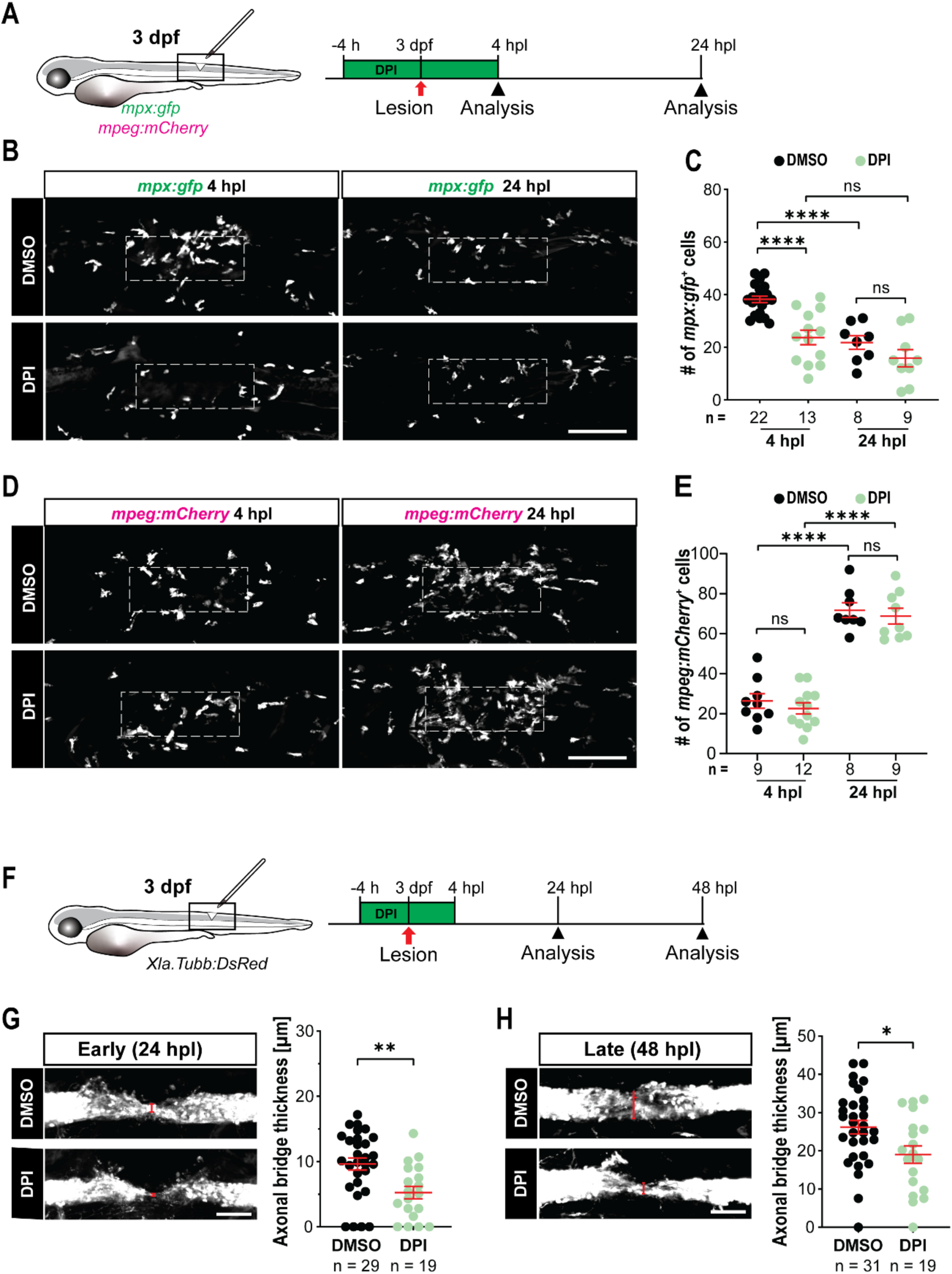
Inhibition of hydrogen peroxide production reduces neutrophil recruitment and impairs axonal regeneration after spinal cord injury. **A:** A schematic timeline of DPI treatment for (B–E) is shown. **B, C:** DPI treatment significantly reduces *mpx:gfp+* neutrophil numbers at 4 hpl but not at 24 hpl compared with DMSO controls. Dashed boxes indicating the quantification area. (Two-way ANOVA, interaction (*F* (1,48) = 3.224, *p* = 0.0789), time (*F* (1,48) = 26.34, *p* < 0.0001), and treatment (*F* (1,48) = 18.61, *p* < 0.0001). Tukey’s multiple comparisons test: *\*\*\*\*p* < 0.0001; ns indicates no significance). **D, E:** DPI treatment does not alter *mpeg:mCherry+* macrophage numbers at 4 and 24 hpl compared with DMSO controls. (Two-way ANOVA, interaction (*F* (1,34) = 0.01227, *p* = 0.9124), time (*F* (1,34) = 170.3, *p* < 0.0001), and treatment (*F* (1,34) = 0.9169, *p* = 0.3451). Tukey’s multiple comparisons test: *\*\*\*\*p* < 0.0001; ns indicates no significance). **F:** A schematic timeline of axonal regeneration analysis for (G, H) is shown. **G, H:** DPI treatment significantly reduces axonal bridge thickness compared with DMSO controls at 24 and 48 hpl. (Two-tailed unpaired t-test: G: *\*p* = 0.0027. H: *\*p* = 0.0178). Error bars show SEM. Scale bar, 100 μm (B, D), 50 μm (G).

**Suppl. Fig. 4.**
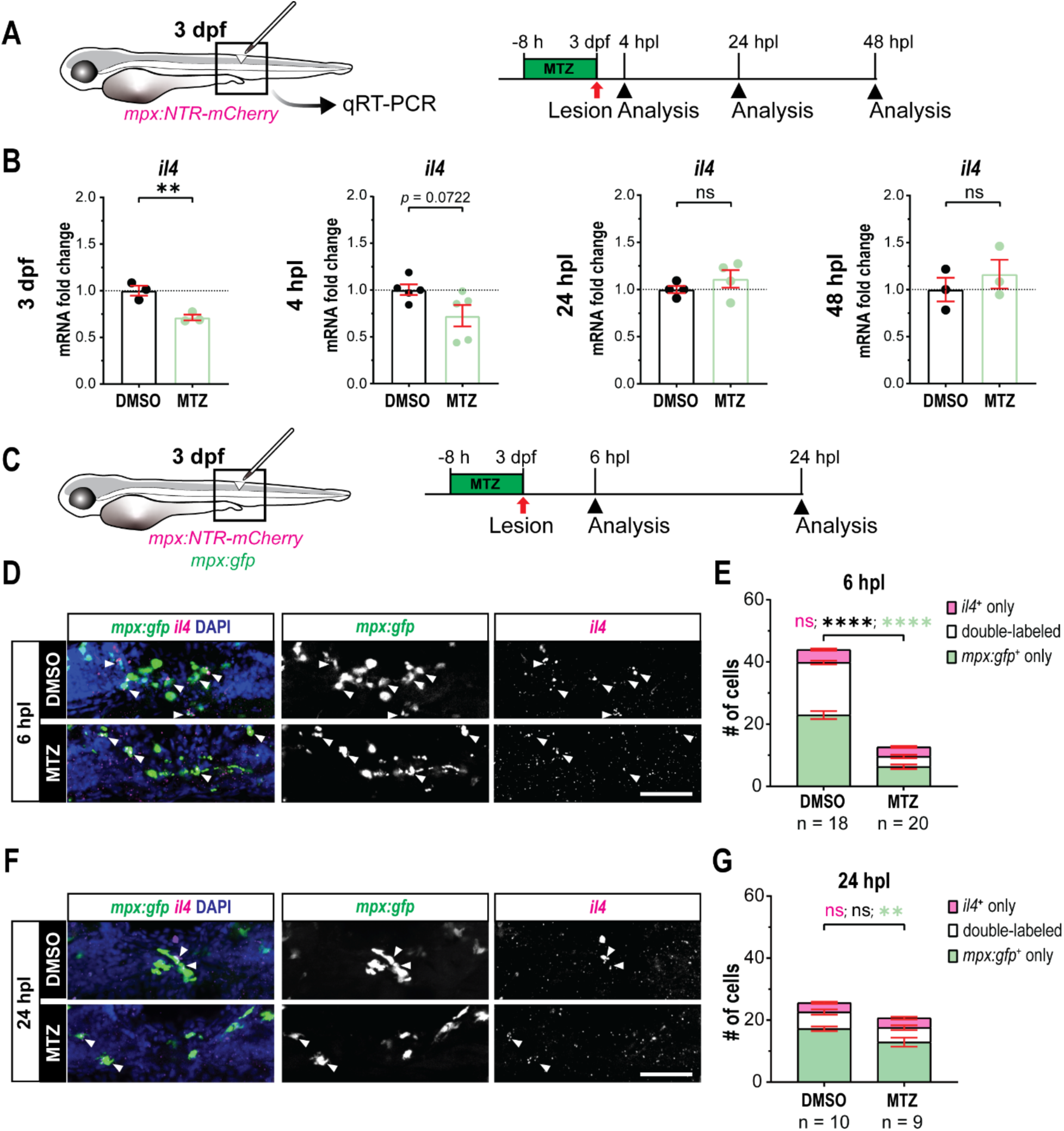
Neutrophil ablation transiently reduces *il4* mRNA abundance but does not detectably alter mRNA levels of other cytokines in unlesioned larvae. **A:** A schematic timeline of qRT-PCR analysis after neutrophil ablation for (B, C) is shown. **B:** Neutrophils ablation reduces the *il4* expression at 3 dpf, but is not detectably changed after lesion. (Two-tailed unpaired t-test: *\*\*p* = 0.0161 at 3 dpl, *p* = 0.0722 at 4 hpl, ns indicates no significance). **C:** A schematic timeline of *il4* expression analysis by HCR after neutrophil ablation for (D–G). **D, E:** Neutrophil ablation significantly reduces the number of *il4*-expressing neutrophils (*il4*⁺, *mpx:gfp*⁺; double-labeled) at 6 hpl. (Two-way ANOVA, interaction (*F* (2,107) = 68.67, *p* < 0.0001), treatment (*F* (1,107) = 332.2, *p* < 0.0001), and cell category (*F* (2,107) = 124.3, *p* < 0.0001). Tukey’s multiple comparisons test: *\*\*\*\*p* < 0.0001. ns indicates no significance). **F, G:** No significant difference is observed in the number of *il4*-expressing neutrophils (*il4*⁺, *mpx:gfp*⁺; double-labeled) following neutrophil ablation at 24 hpl. (Two-way ANOVA, interaction (*F* (2,51) = 4.013, *p* = 0.0241), treatment (*F* (1,51) = 5.907, *p* = 0.0186), and cell category (*F* (2,51) = 120.3, *p* < 0.0001). Tukey’s multiple comparisons test: *\*\*p* = 0.0071. ns indicates no significance). Arrowheads indicate *il4*⁺ neutrophils (*il4*⁺, *mpx:gfp*⁺). Scale bar, 50 μm (D, F). Error bars show SEM.

**Suppl. Fig. 5.**
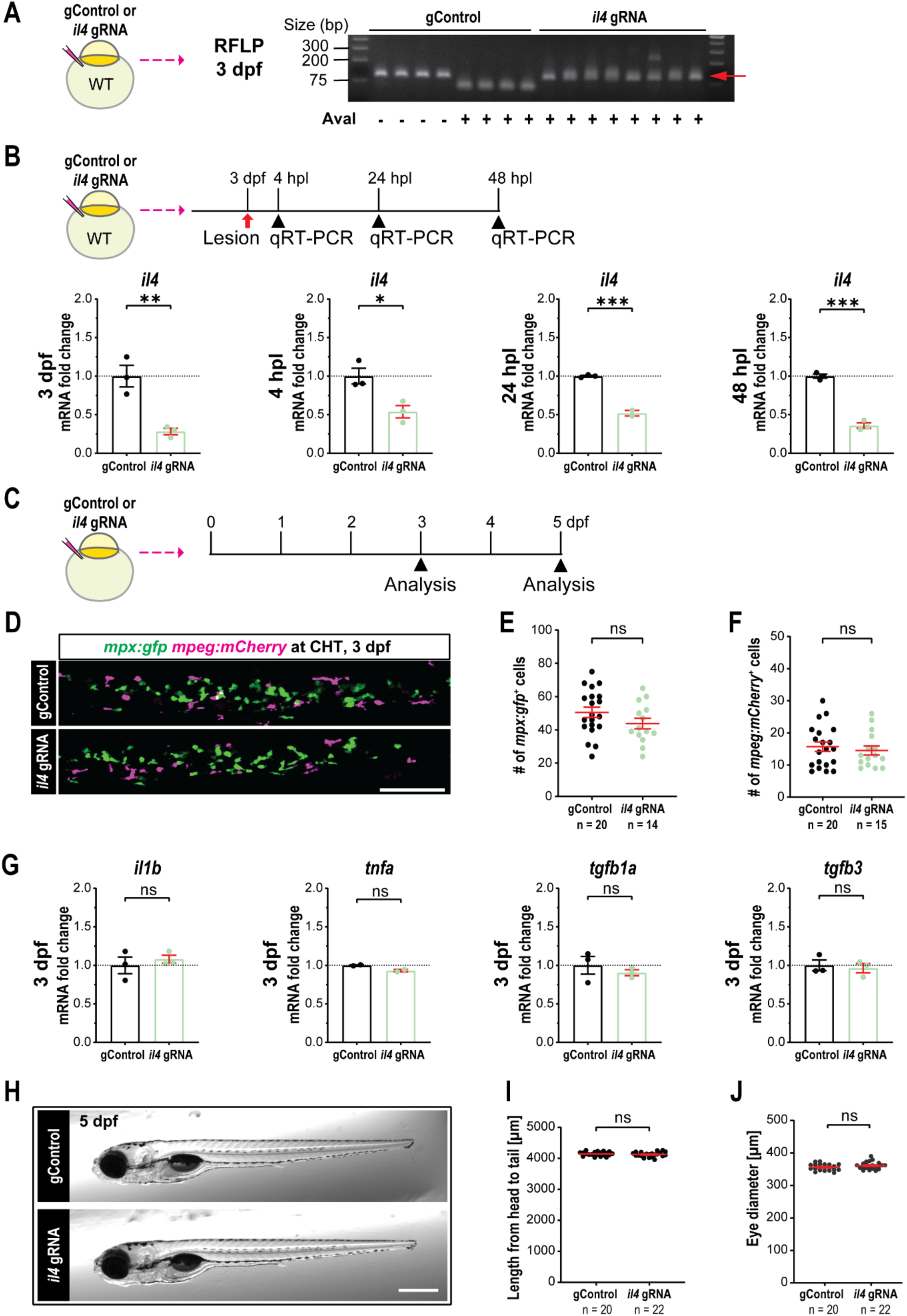
Somatic mutation of *il4* is effective but does not detectably affect immune cell development and larval growth. **A:** RFLP analysis shows effective disruption of genomic DNA after *il4* gRNA injection by resistance to restriction digestion site of the targeted region (red arrows). Each lane represents one larva at 3 dpf. **B:** A qRT-PCR analysis shows *il4* mRNA expression is reduced by *il4 gRNA* injection compared to controls at multiple time points. (Two-tailed unpaired t-test: *\*\*p* = 0.0078 at 3 dpf, *\*p* = 0.0238 at 4 hpl, *\*\*\*p* = 0.0005 at 24 hpl, *\*\*\*p* = 0.0001 at 48 hpl). **C:** A schematic timeline of morphological analysis in *il4 gRNA* injected larvae, corresponding to (D–J), is shown. **D, E, F:** No difference of the developmental number of neutrophils (*mpx:gfp*, green) and macrophages (*mpeg:mCherry*, magenta) was observed in the caudal hematopoietic tissue (CHT) between gControl- and *il4* gRNA-injected larvae at 3 dpf. (Two-tailed unpaired t-test, ns indicates no significance). **J:** A qRT-PCR analysis indicates that *il1b, tnfa*, *tgfb1a* and *tgfb3* expression at 3 dpf are not affect by *il4* gRNA injection, compared to gControl group. (Two-tailed unpaired t-test, ns indicates no significance). **G, H, I:** Overall morphology of gControl- and *il4 gRNA*-injected larvae reveals no difference in body length (E) or eye diameter (F) at 5 dpf. (Two-tailed unpaired t-test, ns indicates no significance). Error bars show SEM. Scale bar, 100 μm (D), 500 μm (H).

**Suppl. Fig. 6.**
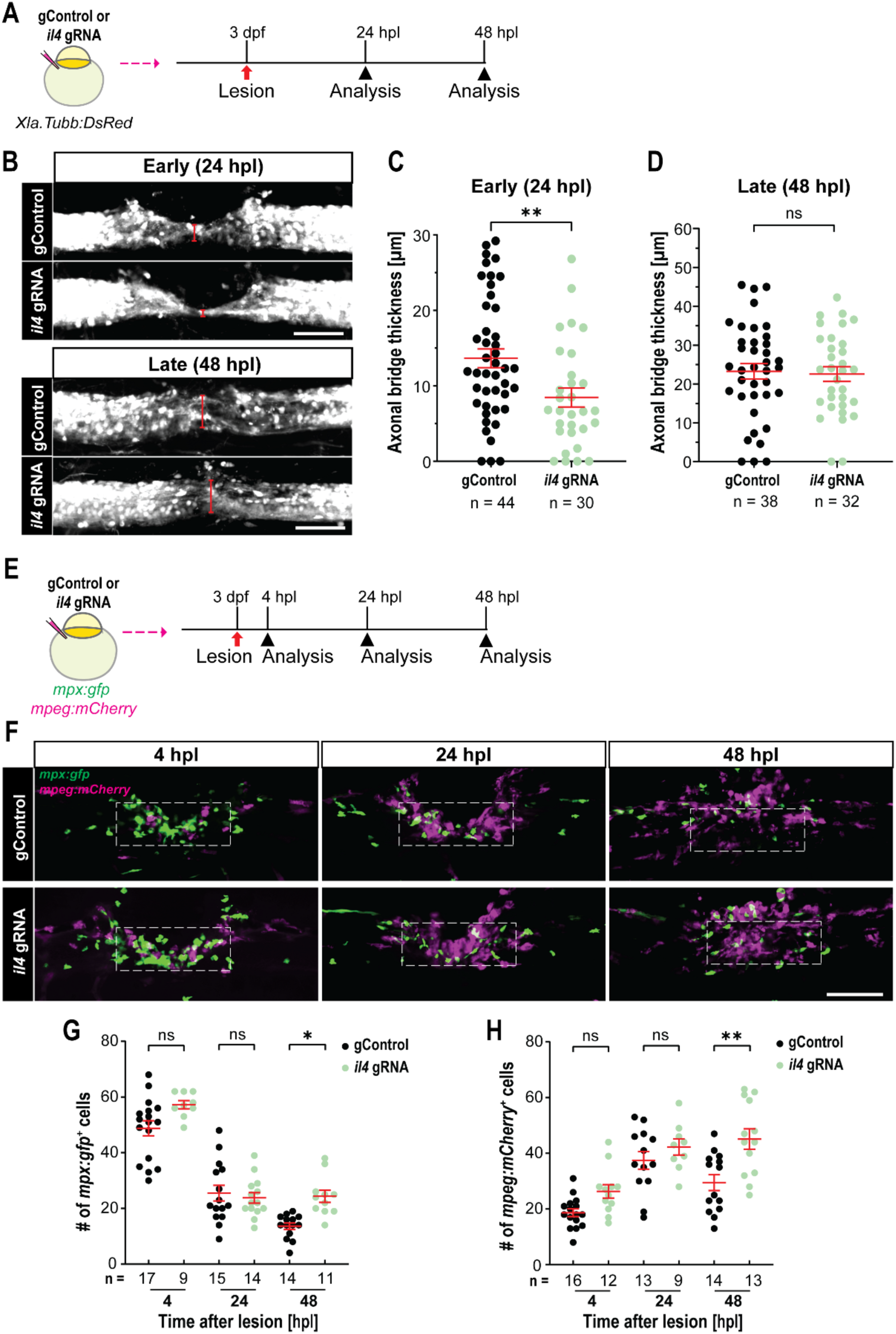
Disruption of *il4* delays axon regrowth and resolution of inflammation. **A:** A schematic timeline of axonal regeneration analysis for (B–D) is shown. **B, C, D:** *Il4* gRNA injection reduces thickness of the axonal bridge at 24 hpl (C), but not at 48 hpl (D). (Two-tailed unpaired t-test: C: *\*\*p* = 0.0058, D: ns indicates no significance). **E:** A schematic timeline of the number of immune cells analysis for (F–H) is shown. **F-H:** *Il4 gRNA* injection slightly increases neutrophil (*mpx:gfp*, green) numbers and macrophage (*mpeg:mCherry*, magenta) numbers at 48 hpl, compared to gControl larvae. Dashed boxes indicate the quantification area. (G: Two-way ANOVA, interaction (*F* (2,74) = 4.225, *p* = 0.0183), time (*F* (2,74) = 114.7, *p* < 0.0001), and treatment (*F* (1,74) = 9.283, *p* = 0.0032). Tukey’s multiple comparisons test: *\*p* = 0.0254; H: Two-way ANOVA, interaction (*F* (2,71) = 1.996, *p* = 0.1434), time (*F* (2,71) = 22.95, *p* < 0.0001), and treatment (*F* (1,71) = 16.56, *p* = 0.0001). Tukey’s multiple comparisons test: *\*\*p* = 0.0015. ns indicates no significance). Error bars show SEM. Scale bar, 50 μm (B), 100 μm (F).

**Suppl. Fig. 7.**
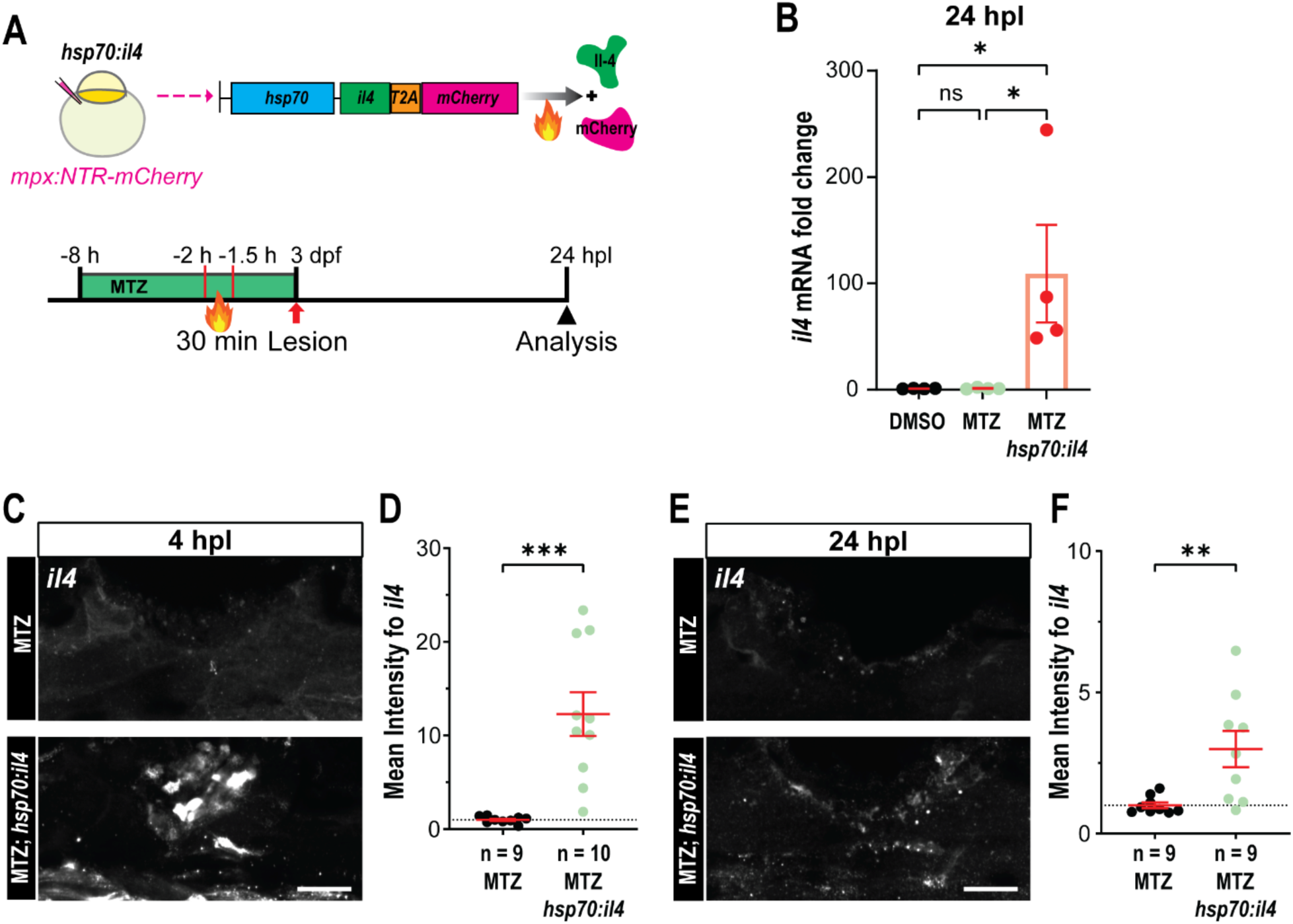
Heat shock–induced *il4* expression increases *il4* mRNA levels in neutrophil-ablated larvae. **A:** A schematic indicating the *hsp70:il4* plasmid construct and experimental timeline for (B-F) is shown. **B:** A qRT-PCR analysis reveals heat shock–induced *il4* overexpression in *hsp70:il4* injected larvae strongly increased *il4* mRNA expression at 24 hpl, compared to controls. (One-way ANOVA, *p* = 0.0271. Tukey’s multiple comparisons test: DMSO vs MTZ *hsp70:il4* (*\*p* = 0.0433), MTZ vs MTZ *hsp70:il4* (*\*p* = 0.0437), ns indicates no significance). **C-F:** Heat shock–induced *il4* expression increases *il4* mRNA signal intensity in neutrophil-ablated larvae with *hsp70:il4* plasmid at 4 hpl (C) and 24 hpl (E). *il4* mRNA was detected by HCR in MTZ-treated only, MTZ-treated and *hsp70:il4* injected larvae after heat shock at 4 hpl (B) and 24 hpl (D). (Two-tailed unpaired t-test: *\*\*\*p* = 0.0009 (C), *\*\*p* = 0.004 (E)). Scale bar, 50 μm. Error bars show SEM.

## Supplementary tables

**Supplementary Table 1.**
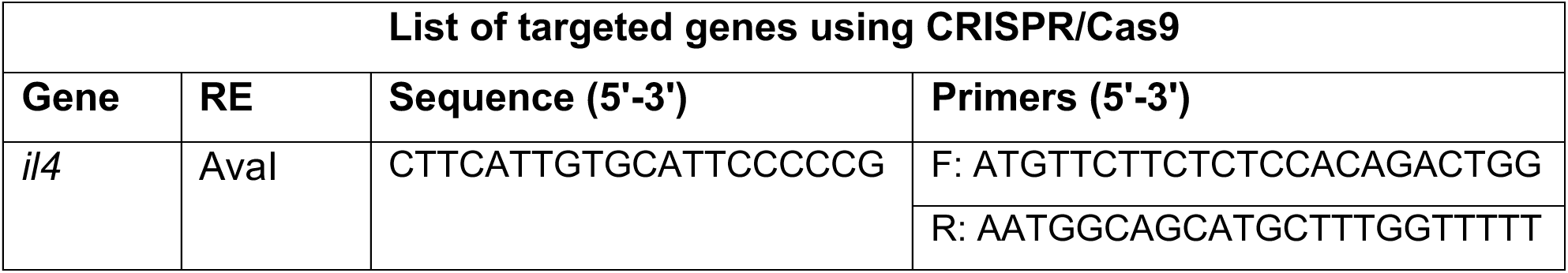
List of targeted genes using CRISPR/Cas9.

**Supplementary Table 2.**
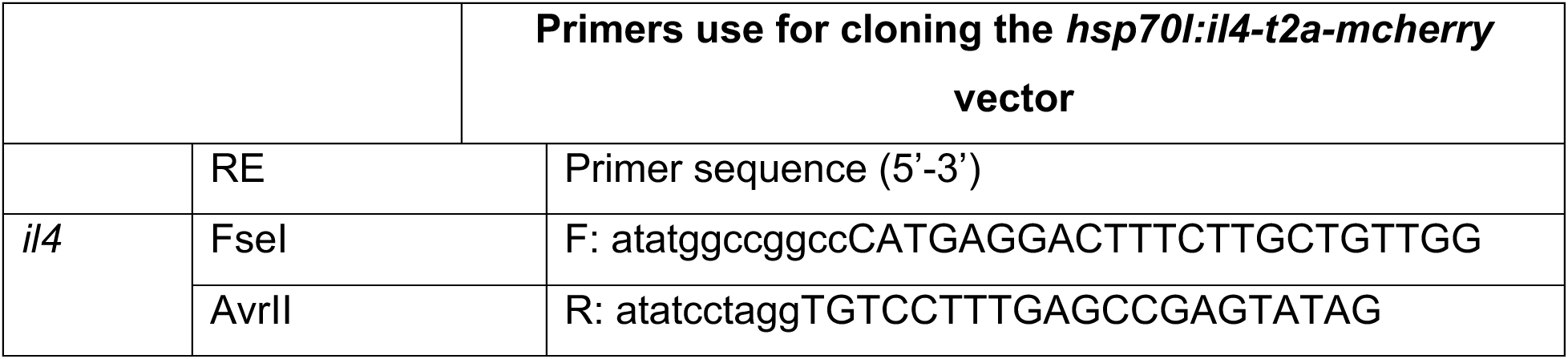
Primers use for cloning the *hsp70l:il4-t2a-mcherry* vector.

**Supplementary Table 3.**
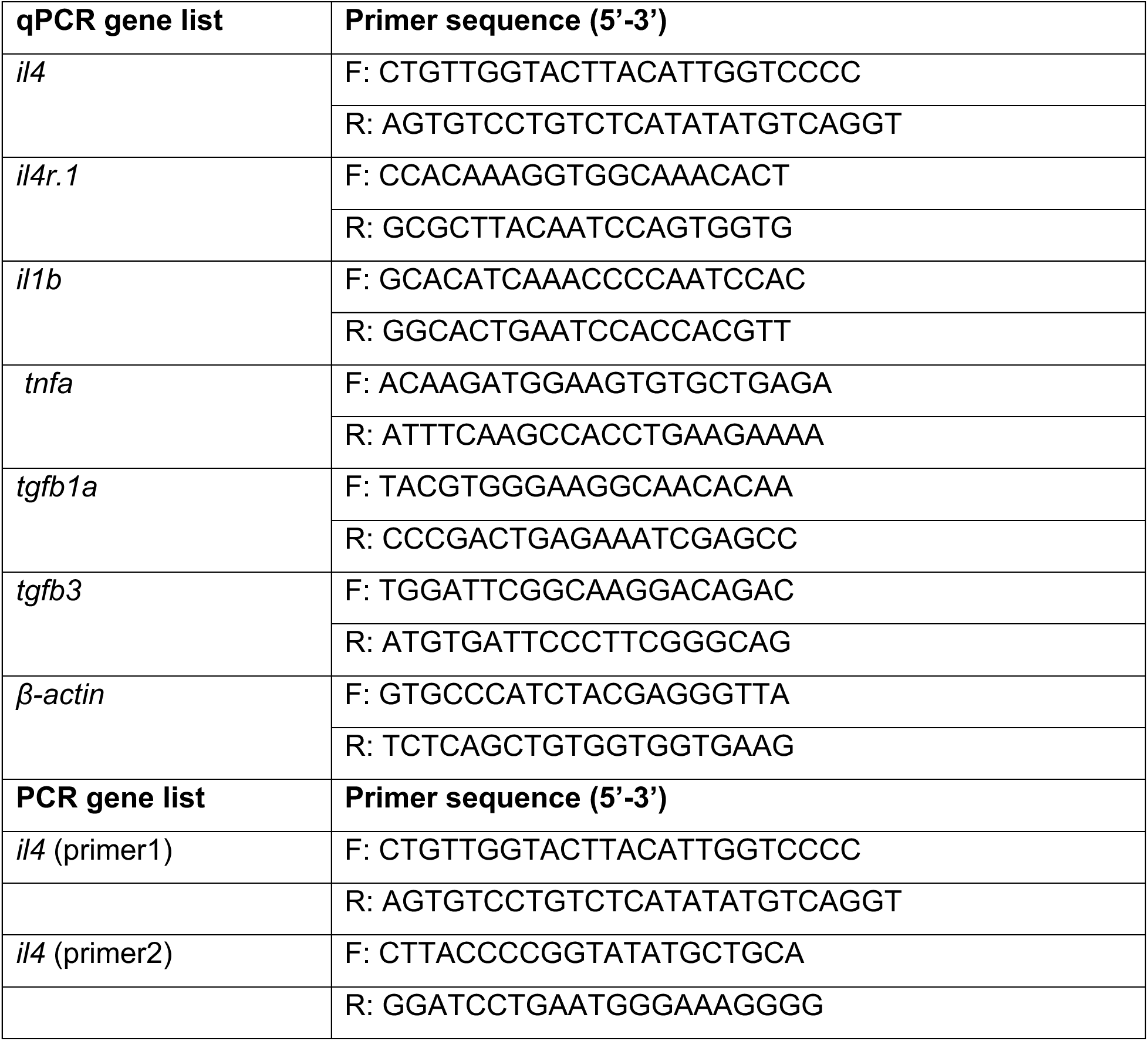
qPCR and PCR primer list.

## CITED LITERATURE

Alper SR, Dorsky RI (2022) Unique advantages of zebrafish larvae as a model for spinal cord regeneration. Front Mol Neurosci 15: 983336

Beck KD, Nguyen HX, Galvan MD, Salazar DL, Woodruff TM, Anderson AJ (2010) Quantitative analysis of cellular inflammation after traumatic spinal cord injury: evidence for a multiphasic inflammatory response in the acute to chronic environment. Brain 133: 433–447

Becker T, Becker CG (2014) Axonal regeneration in zebrafish. Curr Opin Neurobiol 27C: 186-191

Becker T, Becker CG (2022) Regenerative neurogenesis: the integration of developmental, physiological and immune signals. Development 149

Bhattarai P, Thomas AK, Cosacak MI, Papadimitriou C, Mashkaryan V, Froc C, Reinhardt S, Kurth T, Dahl A, Zhang Y et al. (2016) IL4/STAT6 Signaling Activates Neural Stem Cell Proliferation and Neurogenesis upon Amyloid-beta42 Aggregation in Adult Zebrafish Brain. Cell Rep 17: 941–948

Bludau O, Weber A, Bosak V, Kuscha V, Dietrich K, Hans S, Brand M (2024) Inflammation is a critical factor for successful regeneration of the adult zebrafish retina in response to diffuse light lesion. Front Cell Dev Biol 12: 1332347

Bottiglione F, Dee CT, Lea R, Zeef LAH, Badrock AP, Wane M, Bugeon L, Dallman MJ, Allen JE, Hurlstone AFL (2020) Zebrafish IL-4-like Cytokines and IL-10 Suppress Inflammation but Only IL-10 Is Essential for Gill Homeostasis. J Immunol 205: 994–1008

Brand M, Granato M, Nüsslein-Volhard C (2002) Keeping and raising zebrafish. In: Zebrafish: a practical approach, Nüsslein-Volhard C., Dahm R. (eds.)Oxford University Press: Oxford, UK

Brand M, Heisenberg CP, Jiang YJ, Beuchle D, Lun K, Furutani-Seiki M, Granato M, Haffter P, Hammerschmidt M, Kane DA et al. (1996) Mutations in zebrafish genes affecting the formation of the boundary between midbrain and hindbrain. Development 123: 179–190

Brandt E, Woerly G, Younes AB, Loiseau S, Capron M (2000) IL-4 production by human polymorphonuclear neutrophils. J Leukoc Biol 68: 125–130

Cavone L, McCann T, Drake LK, Aguzzi EA, Oprişoreanu AM, Pedersen E, Sandi S, Selvarajah J, Tsarouchas TM, Wehner D et al. (2021) A unique macrophage subpopulation signals directly to progenitor cells to promote regenerative neurogenesis in the zebrafish spinal cord. Dev Cell 56: 1617–1630.e1616

Chen X, Zhang J, Song Y, Yang P, Yang Y, Huang Z, Wang K (2020) Deficiency of anti-inflammatory cytokine IL-4 leads to neural hyperexcitability and aggravates cerebral ischemia-reperfusion injury. Acta Pharm Sin B 10: 1634–1645

Choi HMT, Schwarzkopf M, Fornace ME, Acharya A, Artavanis G, Stegmaier J, Cunha A, Pierce NA (2018) Third-generation in situ hybridization chain reaction: multiplexed, quantitative, sensitive, versatile, robust. Development 145

Concordet JP, Haeussler M (2018) CRISPOR: intuitive guide selection for CRISPR/Cas9 genome editing experiments and screens. Nucleic Acids Res 46: W242–W245

Courtine G, Sofroniew MV (2019) Spinal cord repair: advances in biology and technology. Nat Med 25: 898–908

Daseke MJ, 2nd, Tenkorang-Impraim MAA, Ma Y, Chalise U, Konfrst SR, Garrett MR, DeLeon-Pennell KY, Lindsey ML (2020) Exogenous IL-4 shuts off pro-inflammation in neutrophils while stimulating anti-inflammation in macrophages to induce neutrophil phagocytosis following myocardial infarction. J Mol Cell Cardiol 145: 112-121

Davison JM, Akitake CM, Goll MG, Rhee JM, Gosse N, Baier H, Halpern ME, Leach SD, Parsons MJ (2007) Transactivation from Gal4-VP16 transgenic insertions for tissue-specific cell labeling and ablation in zebrafish. Dev Biol 304: 811–824

de Sena-Tomas C, Rebola Lameira L, Rebocho da Costa M, Naique Taborda P, Laborde A, Orger M, de Oliveira S, Saude L (2024) Neutrophil immune profile guides spinal cord regeneration in zebrafish. Brain Behav Immun 120: 514–531

Denans N, Tran NTT, Swall ME, Diaz DC, Blanck J, Piotrowski T (2022) An anti-inflammatory activation sequence governs macrophage transcriptional dynamics during tissue injury in zebrafish. Nat Commun 13: 5356

Diao Y, Hao M, Xie M, Hu X, Tan R, Wang Z, Rong H, Zhu T (2025) CD47-blocking antibody interferes with neutrophil extracellular traps formation after spinal cord injury to reduce spinal cord edema. J Neuroimmunol 400: 578553

Docampo-Seara A, Cosacack MI, Heilemann K, Kessel F, Oprisoreanu AM, Cark Ö, Zöller D, Arnold J, Bretschneider A, Hnatiuk A et al. (2024) Macrophage crosstalk with neural progenitors and fibroblasts controls regenerative neurogenesis via Sema4ab after spinal cord injury in zebrafish. BioRxiv doi: 10.1101/2024.10.16.618445

Docampo-Seara A, Cosacak MI, Heilemann K, Kessel F, Oprişoreanu A-M, Westphal M, Çark Ö, Zöller D, Arnold J, Bretschneider A et al. (2025) Microglia attenuate regenerative neurogenesis via *sema4ab* after spinal cord injury in zebrafish. bioRxiv: 2024.2010.2016.618445

Dolma S, Kumar H (2021) Neutrophil, Extracellular Matrix Components, and Their Interlinked Action in Promoting Secondary Pathogenesis After Spinal Cord Injury. Mol Neurobiol 58: 4652–4665

Ellett F, Pase L, Hayman JW, Andrianopoulos A, Lieschke GJ (2011) mpeg1 promoter transgenes direct macrophage-lineage expression in zebrafish. Blood 117: e49–56

Fehlings MG, Wilson JR, Harrop JS, Kwon BK, Tetreault LA, Arnold PM, Singh JM, Hawryluk G, Dettori JR (2017) Efficacy and Safety of Methylprednisolone Sodium Succinate in Acute Spinal Cord Injury: A Systematic Review. Global Spine J 7: 116s–137s

Fenn AM, Hall JC, Gensel JC, Popovich PG, Godbout JP (2014) IL-4 signaling drives a unique arginase+/IL-1beta+ microglia phenotype and recruits macrophages to the inflammatory CNS: consequences of age-related deficits in IL-4Ralpha after traumatic spinal cord injury. J Neurosci 34: 8904–8917

Flanagan-Steet H, Fox MA, Meyer D, Sanes JR (2005) Neuromuscular synapses can form in vivo by incorporation of initially aneural postsynaptic specializations. Development 132: 4471–4481

Francos-Quijorna I, Amo-Aparicio J, Martinez-Muriana A, Lopez-Vales R (2016) IL-4 drives microglia and macrophages toward a phenotype conducive for tissue repair and functional recovery after spinal cord injury. Glia 64: 2079–2092

Galli SJ, Borregaard N, Wynn TA (2011) Phenotypic and functional plasticity of cells of innate immunity: macrophages, mast cells and neutrophils. Nat Immunol 12: 1035–1044

Gideon HP, Phuah J, Junecko BA, Mattila JT (2019) Neutrophils express pro- and anti-inflammatory cytokines in granulomas from Mycobacterium tuberculosis-infected cynomolgus macaques. Mucosal Immunol 12: 1370–1381

Greenhalgh AD, David S, Bennett FC (2020) Immune cell regulation of glia during CNS injury and disease. Nat Rev Neurosci 21: 139–152

Guedes JR, Ferreira PA, Costa J, Laranjo M, Pinto MJ, Reis T, Cardoso AM, Lebre C, Casquinha M, Gomes M et al. (2023) IL-4 shapes microglia-dependent pruning of the cerebellum during postnatal development. Neuron 111: 3435–3449 e3438

John N, Fleming T, Kolb J, Lyraki O, Vasquez-Sepulveda S, Parmar A, Kim K, Tarczewska M, Gupta P, Singh K et al. (2025) Biphasic inflammation control by fibroblasts enables spinal cord regeneration in zebrafish. Cell Rep 44: 116469

John N, Kolb J, Wehner D (2022) Mechanical spinal cord transection in larval zebrafish and subsequent whole-mount histological processing. STAR Protoc 3: 101093

Keatinge M, Tsarouchas TM, Munir T, Porter NJ, Larraz J, Gianni D, Tsai HH, Becker CG, Lyons DA, Becker T (2021) CRISPR gRNA phenotypic screening in zebrafish reveals pro-regenerative genes in spinal cord injury. PLoS Genet 17: e1009515

Kitade K, Kobayakawa K, Saiwai H, Matsumoto Y, Kawaguchi K, Iida K, Kijima K, Iura H, Tamaru T, Haruta Y et al. (2023) Reduced Neuroinflammation Via Astrocytes and Neutrophils Promotes Regeneration After Spinal Cord Injury in Neonatal Mice. J Neurotrauma 40: 2566–2579

Kumar H, Choi H, Jo MJ, Joshi HP, Muttigi M, Bonanomi D, Kim SB, Ban E, Kim A, Lee SH et al. (2018) Neutrophil elastase inhibition effectively rescued angiopoietin-1 decrease and inhibits glial scar after spinal cord injury. Acta neuropathologica communications 6: 73

Kurimoto T, Yin Y, Habboub G, Gilbert HY, Li Y, Nakao S, Hafezi-Moghadam A, Benowitz LI (2013) Neutrophils express oncomodulin and promote optic nerve regeneration. J Neurosci 33: 14816–14824

Liebold I, Al Jawazneh A, Casar C, Lanzloth C, Leyk S, Hamley M, Wong MN, Kylies D, Grafe SK, Edenhofer I et al. (2024) Apoptotic cell identity induces distinct functional responses to IL-4 in efferocytic macrophages. Science 384: eabo7027

Mirat O, Sternberg JR, Severi KE, Wyart C (2013) ZebraZoom: an automated program for high-throughput behavioral analysis and categorization. Front Neural Circuits 7: 107

Montgomery DC (2017) Design and Analysis of Experiments. John Wiley & Sons, Hoboken, NJ

Moreno-Loaiza O, Soares VC, de Assumpcao Souza M, Vera-Nunez N, Rodriguez de Yurre Guirao A, da Silva TP, Pozes AB, Perticarrari L, Monteiro E, Albino MC et al. (2025) IL-1beta enhances susceptibility to atrial fibrillation in mice by acting through resident macrophages and promoting caspase-1 expression. Nat Cardiovasc Res 4: 312-329

Moreno-Mateos MA, Vejnar CE, Beaudoin JD, Fernandez JP, Mis EK, Khokha MK, Giraldez AJ (2015) CRISPRscan: designing highly efficient sgRNAs for CRISPR-Cas9 targeting in vivo. Nat Methods 12: 982–988

Nguyen HX, Hooshmand MJ, Saiwai H, Maddox J, Salehi A, Lakatos A, Nishi RA, Salazar D, Uchida N, Anderson AJ (2017) Systemic Neutrophil Depletion Modulates the Migration and Fate of Transplanted Human Neural Stem Cells to Rescue Functional Repair. J Neurosci 37: 9269–9287

Niethammer P, Grabher C, Look AT, Mitchison TJ (2009) A tissue-scale gradient of hydrogen peroxide mediates rapid wound detection in zebrafish. Nature 459: 996–999

Nüsslein-Volhard CD, Ralf (2002) Zebrafish: a practical approach. Oxford University Press, New York

Ohnmacht J, Yang Y, Maurer GW, Barreiro-Iglesias A, Tsarouchas TM, Wehner D, Sieger D, Becker CG, Becker T (2016) Spinal motor neurons are regenerated after mechanical lesion and genetic ablation in larval zebrafish. Development 143: 1464–1474

Oprişoreanu AM, Ryan F, Richmond C, Dzekhtsiarova Y, Carragher NO, Becker T, David S, Becker CG (2023) Drug screening in zebrafish larvae reveals inflammation-related modulators of secondary damage after spinal cord injury in mice. Theranostics https://www.thno.org/v13p2531

Peri F, Nusslein-Volhard C (2008) Live imaging of neuronal degradation by microglia reveals a role for v0-ATPase a1 in phagosomal fusion in vivo. Cell 133: 916–927

Renshaw SA, Loynes CA, Trushell DM, Elworthy S, Ingham PW, Whyte MK (2006) A transgenic zebrafish model of neutrophilic inflammation. Blood 108: 3976–3978

Richter M, Diesterbeck E, Pylaeva E, Labusek N, Koster C, Nagel D, Karsch L, Fischer AJ, Sous M, Jung M et al. (2025) Hypoxic-ischemic brain injury in neonatal mice sequentially recruits neutrophils with dichotomous phenotype and function. Nat Commun 16: 9696

Robertson AL, Holmes GR, Bojarczuk AN, Burgon J, Loynes CA, Chimen M, Sawtell AK, Hamza B, Willson J, Walmsley SR et al. (2014) A zebrafish compound screen reveals modulation of neutrophil reverse migration as an anti-inflammatory mechanism. Sci Transl Med 6: 225ra229

Rodgers KA, Kigerl KA, Schwab JM, Popovich PG (2022) Immune dysfunction after spinal cord injury - A review of autonomic and neuroendocrine mechanisms. Curr Opin Pharmacol 64: 102230

Rueden CT, Schindelin J, Hiner MC, DeZonia BE, Walter AE, Arena ET, Eliceiri KW (2017) ImageJ2: ImageJ for the next generation of scientific image data. BMC Bioinformatics 18: 529

Saiwai H, Ohkawa Y, Yamada H, Kumamaru H, Harada A, Okano H, Yokomizo T, Iwamoto Y, Okada S (2010) The LTB4-BLT1 axis mediates neutrophil infiltration and secondary injury in experimental spinal cord injury. Am J Pathol 176: 2352–2366

Sas AR, Carbajal KS, Jerome AD, Menon R, Yoon C, Kalinski AL, Giger RJ, Segal BM (2020) A new neutrophil subset promotes CNS neuron survival and axon regeneration. Nat Immunol 21: 1496–1505

Schindelin J, Arganda-Carreras I, Frise E, Kaynig V, Longair M, Pietzsch T, Preibisch S, Rueden C, Saalfeld S, Schmid B et al. (2012) Fiji: an open-source platform for biological-image analysis. Nat Methods 9: 676–682

Schwarzkopf M, Liu MC, Schulte SJ, Ives R, Husain N, Choi HMT, Pierce NA (2021) Hybridization chain reaction enables a unified approach to multiplexed, quantitative, high-resolution immunohistochemistry and in situ hybridization. Development 148

Stewart AN, Bosse-Joseph CC, Kumari R, Bailey WM, Park KA, Slone VK, Gensel JC (2025) Nonresolving Neuroinflammation Regulates Axon Regeneration in Chronic Spinal Cord Injury. J Neurosci 45

Tendolkar A, Mokalled MH (2025) Mechanisms underpinning spontaneous spinal cord regeneration. Development 152

Tsarouchas TM, Wehner D, Cavone L, Munir T, Keatinge M, Lambertus M, Underhill A, Barrett T, Kassapis E, Ogryzko N et al. (2018) Dynamic control of proinflammatory cytokines Il-1β and Tnf-α by macrophages in zebrafish spinal cord regeneration. Nat Commun 9: 4670

Tsata V, Möllmert S, Schweitzer C, Kolb J, Möckel C, Böhm B, Rosso G, Lange C, Lesche M, Hammer J et al. (2021) A switch in pdgfrb(+) cell-derived ECM composition prevents inhibitory scarring and promotes axon regeneration in the zebrafish spinal cord. Dev Cell 56: 509–524.e509

Tsata V, Wehner D (2021) Know How to Regrow-Axon Regeneration in the Zebrafish Spinal Cord. Cells 10

Wehner D, Tsarouchas TM, Michael A, Haase C, Weidinger G, Reimer MM, Becker T, Becker CG (2017) Wnt signaling controls pro-regenerative Collagen XII in functional spinal cord regeneration in zebrafish. Nat Commun 8: 126

Westerfield M (2007) The Zebrafish Book. A Guide for the Laboratory Use of Zebrafish (Danio rerio). University of Oregon Press, Eugene

Yates AG, Jogia T, Gillespie ER, Couch Y, Ruitenberg MJ, Anthony DC (2021) Acute IL-1RA treatment suppresses the peripheral and central inflammatory response to spinal cord injury. J Neuroinflammation 18: 15

